# Heterochromatin rewiring and domain disruption-mediated chromatin compaction during erythropoiesis

**DOI:** 10.1101/2021.08.12.456090

**Authors:** Dong Li, Fan Wu, Shuo Zhou, Xiao-Jun Huang, Hsiang-Ying Lee

**Author notes:** These authors contributed equally to this work (Dong Li, Fan Wu).

## Abstract

Development of mammalian red blood cells involves progressive chromatin compaction and subsequent enucleation in terminal stages of differentiation, but the molecular mechanisms underlying the three-dimensional chromatin reorganization and compaction remains obscure. Here, we systematically analyze the distinct features of higher-order chromatin in purified populations of primary human erythroblasts. Our results reveal that while heterochromatin regions undergo substantial compression, select transcription competent regions with active transcription signature are preferentially maintained to achieve a highly-compacted yet functional chromatin state in terminal erythropoiesis, which is about 20-30% of the nuclear volume compared to that of erythroid progenitors. While the partition of euchromatic and heterochromatic regions (compartment A and B) remain mostly unchanged, H3K9me3 marks relocalize to the nuclear periphery and a significant number of H3K9me3 long-range interactions are formed in the three-dimensional rewiring during terminal erythroid chromatin condensation. Moreover, ∼63% of the topologically associating domain (TAD) boundaries are disrupted, while certain TADs with active chromatin modification are selectively maintained during terminal erythropoiesis. The most well-maintained TADs are enriched for chromatin structural factors CTCF and SMC3, as well as factors and marks of the active transcription state. Finally, we demonstrate that the erythroid master regulator GATA1 involves in safeguarding select essential chromatin domains during terminal erythropoiesis. Our study therefore delineate the molecular characteristics of a development-driven chromatin compaction process, which reveals transcription competence as a key determinant of the select domain maintenance to ensure appropriate gene expression during immense chromatin compaction.

## Introduction

Erythropoiesis is defined by a stepwise process of proliferation and differentiation from hematopoietic stem and progenitor cells (HSPCs) to mature red blood cells. Differentiation from the late erythroid progenitor cells, colony-forming units-erythroid (CFU-E), to mature red blood cells, termed terminal erythropoiesis, involves 3-5 cell divisions with morphologically distinguishable erythroblasts which include pro-(Pro-E), basophilic- (Baso-E), polychromatic- (Poly-E), and orthochromatic- erythroblasts (Ortho-E)^1, 2^. During mammalian terminal erythropoiesis, erythroblasts undergo drastic nuclear compaction which is associated with chromatin condensation, followed by extrusion of the pyknotic nucleus in the final stage of terminal differentiation^3–7^.

Chromatin condensation starts from the onset of terminal erythropoiesis. The nuclear volume decreases along with each terminal cell division and results in a final nuclear volume which is several fold smaller than the progenitor stage^4^. Erythroid chromatin condensation is important for terminal erythropoiesis, and defects in the process are observed frequently in myelodysplastic syndromes and megaloblastic anemia^8^. Several factors were shown to be involved in the erythroid chromatin condensation, including regulators of histone acetylation, such as HDACs and GCN5^9–11^. In addition, global DNA demethylation was observed in both human and murine erythropoiesis^12, 13^. However, the molecular characteristics of chromatin condensation still remains elusive. Furthermore, the dynamics and regulation of higher-order chromatin structure during this process is largely unknown.

Three-dimensional organization of chromosomes promotes the long-range chromatin interactions especially between enhancers and promoters that are important for maintaining sophisticated gene regulatory networks in multicellular organisms^14, 15^. Self-associating chromatin domains identified in Hi-C assays, Topologically Associating Domains (TADs), which exhibit high levels of intradomain interactions and low levels of interdomain interactions, facilitate proper gene activation and repression by restricting *cis*-regulatory element interactions with target promoters^16–20^. TAD boundaries are largely conserved among different cell types and even closely related species^18, 19^. At larger scales, interactions between TADs can give rise to two main types of spatial compartments referred to as ‘‘A’’ and ‘‘B’’, which are functionally related to gene activation and repression, respectively and could switch frequently during differentiation^21^.

Heterochromatin plays vital roles in fundamentally important processes such as maintenance of genome stability and control of cell differentiation^22^. A dense layer of heterochromatin can be found underneath the nuclear lamina in most mammalian cells, which has been identified as Lamin-Associated Domains (LADs). The lamina-associated heterochromatin regions are enriched for repressive histone modifications, such as H3K9me2, H3K9me3, and H3K27me3^23^. Proper heterochromatin deposition is essential for packaging of the genome to ensure appropriate gene silencing.

Terminal erythropoiesis is coupled with global down regulation of transcriptional activity in both mouse and human^24–26^, along with gradually diminished chromatin accessibility for most transcription factor binding sites, with the exception for the motifs of erythroid master regulators - GATA1 and KLF1^26^. The accessibility of GATA1 and KLF1 binding motifs may indicate that GATA1 and KLF1 govern transcription till the end of terminal erythropoiesis. Moreover, GATA1 was shown to bookmark mitotic chromatin for rapid reactivation of its target genes upon entry into G1 phase of cell cycle^27^. Nonetheless, it remains unclear whether GATA1 and KLF1 function directly to control terminal erythropoiesis and whether their function involves reorganization of higher-order chromatin architecture while global chromatin condensation occurs.

Here, we demonstrate that the formation of long-range heterochromatin interactions reshapes the three-dimensional chromatin architecture, serving as a major participant of erythroid chromatin condensation. Globally, TAD structures are largely attenuated and TAD boundary strength becomes significantly weakened in terminal erythroid development, but the TADs with active chromatin modification are selectively maintained. Furthermore, our results demonstrate that GATA1 is involved in maintaining the active chromatin topology and terminal erythroid gene expression program during terminal erythropoiesis. Our study reveals a unique chromatin organization process involving 3-5 fold nuclear condensation during human terminal erythropoiesis, which is characterized by *de novo* establishment and reorganization of long-range heterochromatin interactions, as well as selective maintenance of transcription competent chromatin domains enriched for active transcription signature.

## Results

### Heterochromatic chromatin regions undergo more pronounced compaction in the global nuclear condensation of terminal erythropoiesis

To obtain defined terminal erythroid cell populations with high purity, we established an efficient method to isolate cord blood CD34^+^ cell-derived primary human erythroblasts at distinct terminal stages by modifying previously reported methods (Fig. 1a)^28, 29^. The CD71^+^ CD235a^+^ CD36^+^ CD45^low^ population are defined as erythroblasts; within the erythroblast population, the CD117(c-kit)^high^ CD105^high^ cells are proerythroblasts (Pro-Es), CD117^dim^ CD105^dim^ cells are basophilic erythroblasts (Baso-Es), CD117^-^ CD105^-^ FSC^high^ cells are polychromatic erythroblasts (Poly-Es), and CD117^-^ CD105^-^ FSC^low^ cells are orthochromatic erythroblasts (Ortho-Es) (Fig. 1a and Extended Data Fig. 1a). To eliminate the impact of cell cycle, G0/G1 cells were sorted based on Hoechst33342 staining (Extended Data Fig. 1a). Each population showed the morphological characteristics and accumulation of hemoglobin of the respective stages (Fig. 1b and Extended Data Fig. 1b,c). The average nuclear diameter and area decreased from Pro-E to Ortho-E stage (Extended Data Fig. 1d). Interestingly, human terminal erythroblasts have a non-canonical cell cycle distribution which is featured by high enrichment of S-phase cells, especially in Pro-E and Baso-E, and this is consistent with murine terminal erythroblasts (Extended Data Fig. 1e)^30^. To validate whether the purified cell populations can recapitulate normal terminal erythroid development, we cultured the sorted cells *in vitro*. The mitotic capacity of Pro-E and Ortho-E was characterized previously — one Pro-E cell can proliferate three times and give rise to eight Ortho-E cells, while Ortho-Es are unable to further divide^31^. Cell number quantification of the *in vitro* cultured Pro-E, Baso-E, Poly-E, and Ortho-E on Day 1 – Day 4 showed that Pro-E, Baso-E, Poly-E, and Ortho-E underwent three, two, one, and zero times of mitosis, respectively (Extended Data Fig. 1f). After four days in culture, sorted Pro-Es showed decreased CD105, CD117 expression and cell size (FSC), and gave rise to enucleated cells (Extended Data Fig. 1g-j). Taken together, these results validated the identity of purified populations of erythroblasts.

**Fig. 1:**
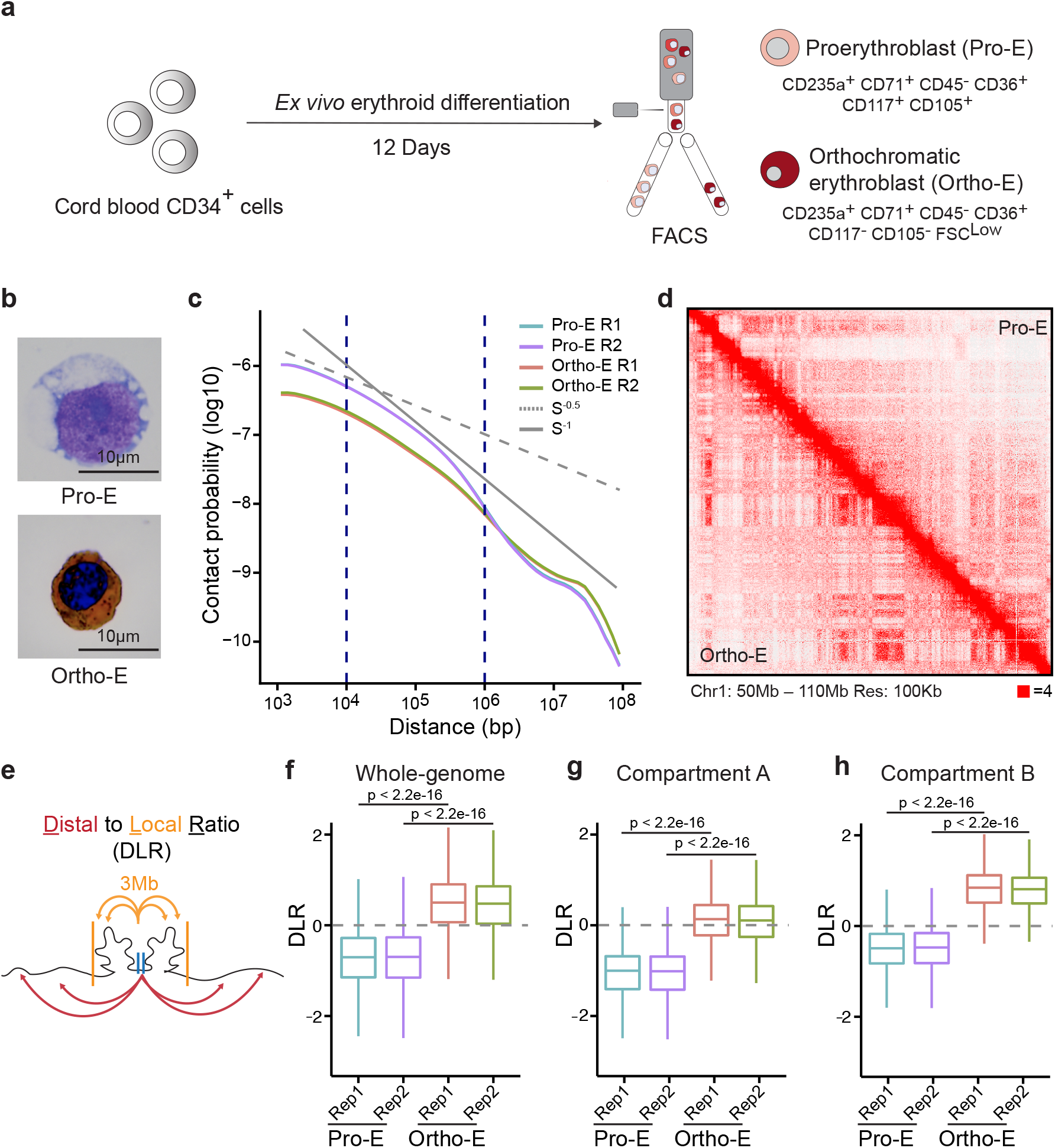
Dynamic chromatin architecture during progressive nuclear condensation of terminal erythropoiesis. **a,** Experimental scheme for isolating primary human erythroblasts of terminal stages via fluorescence-activated cell sorting (FACS). **b,** Representative images of purified primary pro-erythroblast (Pro-E) and orthochromatic erythroblast (Ortho-E) by benzidine-Giemsa staining. **c,** Hi-C contact probability as a function of distance in Pro-E (cyan, purple) and Ortho-E (red, green). P(s)∼S^-0.5^ indicates the mitotic state, ∼S^-1^ indicates the fractal globule state. **d,** Hi-C contact matrices of chromosome 1 (50–110 Mb) in Pro-E and Ortho-E cells. Juicebox parameters were balanced normalized at 100 kb resolution. **e,** Schematic illustration of DLR (Distal to Local Ratio). **f-h,** DLR for chromatin interactions in the (**f**) whole genome, (**g**) compartment A and (**h**) compartment B of Pro-E and Ortho-E.

To investigate the chromatin condensation and architecture changes in terminal erythropoiesis, we conducted *in situ* Hi-C in primary human Pro-Es and Ortho-Es isolated as described above (Fig. 1b). The Hi-C assay generated over half a billion uniquely aligned contacts for each biological replicate which were highly correlated at all resolutions of Pro-E and Ortho-E. The overall maximum resolution is about 10 kb. We found much more frequent long-range intra-chromosome interactions in Ortho-Es, but more short-range intra-chromosome interactions in Pro-Es, based on the contact probability (Fig. 1c,d and Supplementary Table 1). The non-cell-cycle-specific samples of Pro-E and Ortho-E also showed consistent results (Extended Data Fig. 2a). The inter-chromosome interactions also increased from Pro-E to Ortho-E stage, markedly (Supplementary Table 1). We also performed MNase-seq to characterize the short-range chromatin compaction at the nucleosome level. Our results indicate that the nucleosome repeat length (NRL) reduced from Pro-E (∼195 bp) to Orhto-E (∼192 bp) stage, at whole genomic level, in consistent with the overall compaction of chromatin during terminal erythropoiesis (Extended Data Fig. 2b).

Identification of compartments A and B between Pro-E and Ortho-E stages based on principal component 1 (PC1) value showed that over 99% of the genome maintained the same compartment state through terminal erythropoiesis (Extended Data Fig. 2c). To further verify the assignment of A/B compartments, we performed Cleavage Under Targets and Release Using Nuclease (CUT&RUN) of H3K27ac, H3K9me3, and RNA-seq in Pro-E and Ortho-E cells. Results show that A compartments are highly enriched for H3K27ac and highly-expressed genes (Extended Data Fig. 2d,e), while B compartments are highly enriched for H3K9me3 (Extended Data Fig. 2f). We also calculated the compartment strength^32^, and no evident differences were found between Pro-Es and Ortho-Es (Extended Data Fig. 2g). We then analyzed the degree of compaction at the whole-genome level, compartments A and B by calculating the Distal to Local Ratio (DLR) which is defined by Log2 ratio of distal Hi-C interactions (greater than 3Mb) to local Hi-C interactions (within 3 Mb) on the same chromosome, as well as the Inter-Chromosomal Fraction of Interactions (ICF) which refers to the ratio of inter-chromosomal interactions to the total number of interactions (inter- and intra-chromosomal interactions) of a designated 15kb region (Fig. 1e and Extended Data Fig. 2h). At the whole-genome level, the median of DLR increased from -0.70 to 0.49 in Pro-E to Ortho-E transition, demonstrating the occurrence of chromatin compaction. The median of DLR in compartments A increased from -1.14 to 0.17 in Pro-E to Ortho-E transition, while in compartments B, DLR median increased from -0.46 to 0.79 (Fig. 1f-h). The higher DLR in compartments B of Ortho-E compared to that in compartments A of Ortho-E suggests compartments B undergo a higher degree of compaction. Generally, ICF is higher in Ortho-E cells than Pro-E cells, but no obvious difference was observed when comparing ICF among whole-genome, compartments A and B within Pro-E or Ortho-E independently (Extended Data Fig. 2i-k). These results indicate the inactive compartments B exhibit a higher degree of chromatin compaction than the active compartments A.

Taken together, in human terminal erythroid differentiation, the frequency of long-range interactions increased along with the ongoing nuclear compaction, consistent with the overall reduction of nucleosome repeat length. Interestingly, a higher degree of chromatin compaction was found in compartments B, relative to compartments A, but no major compartment switch events were found.

### Chromatin condensation in terminal erythropoiesis is coupled with relocalization and establishment of long-range interactions of heterochromatin

To dissect the underlying mechanisms of higher compaction within compartments B, we next analyzed chromatin interactions of A-A, B-B and A-B, respectively. Plotting the log ratio of observed versus expected contacts revealed a preferential gain of B-B interactions and loss of A-A and A-B interactions during the transition from Pro-E to Ortho-E cells (Fig. 2a). We also validated these results by using an alternative approach based on eigenvector values (Fig. 2b). Contact matrices revealed that Ortho-E cells have a much higher frequency of long-range interactions in compartments B than Pro-E cells, whereas not much difference was observed between compartments A of Pro-E and those of Ortho-E, demonstrating augmented interactions among compartments B (inactive) play a more predominant role than compartments A (active) in terminal erythroid chromatin compaction (Fig. 2c).

**Fig. 2:**
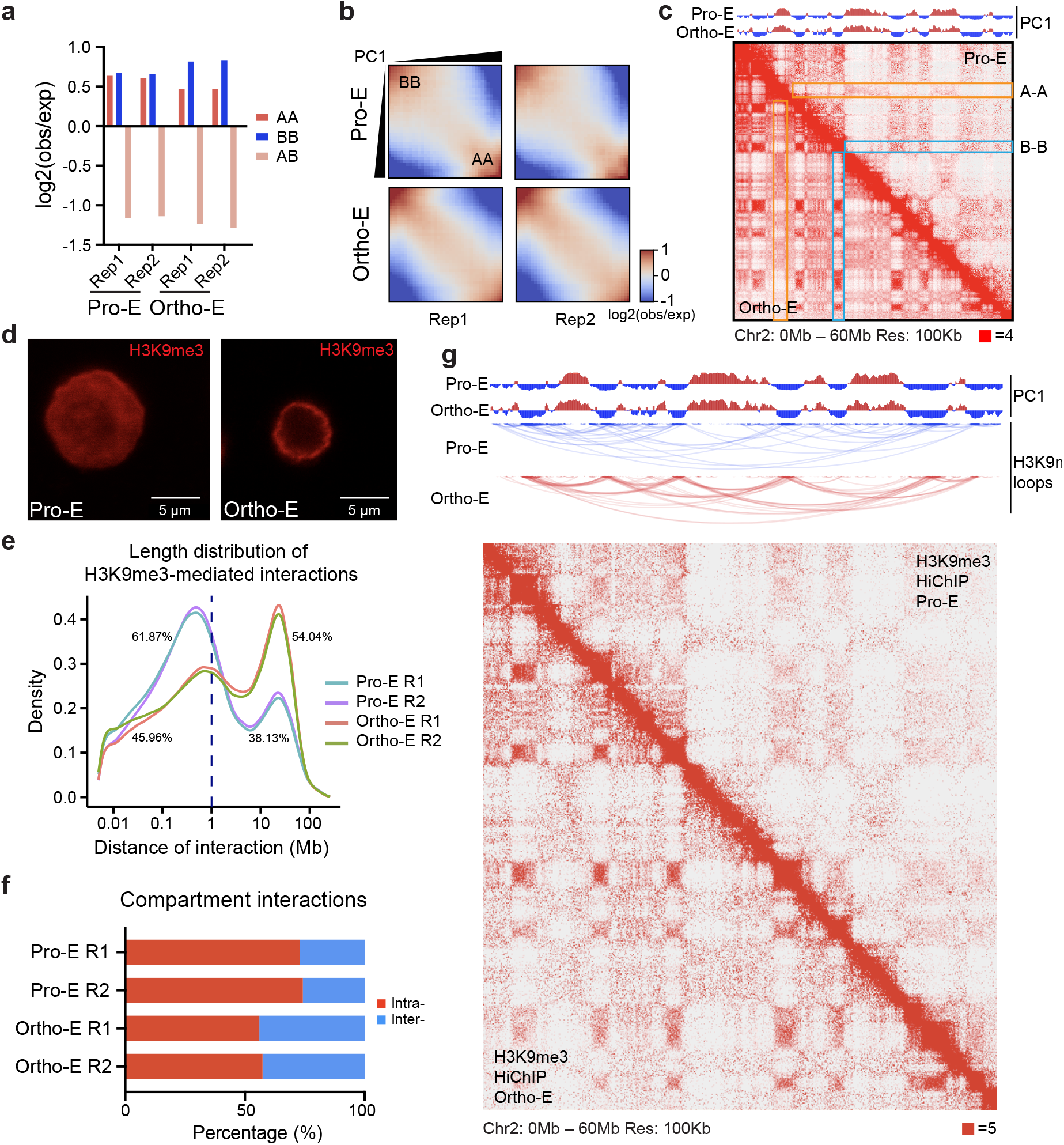
Chromatin condensation in terminal erythropoiesis is coupled with relocalization and establishment of long-range interactions of heterochromatin. **a**, The log2 ratio for observed (obs) versus expected (exp) Hi-C contact intensity values of A-A, B-B and A-B compartment interactions in Pro-E to Ortho-E cells. **b**, Saddle plots showing the compartment interactions change between Pro-E and Ortho-E. **c**, (Top) PC1 value of Hi-C analysis indicating chromatin compartmentalization. Red: compartment A; blue: compartment B. (Bottom) Hi-C contact matrices for chromosome 2 (0–60 Mb) of Pro-E and Ortho-E cells. Juicebox parameters were balanced normalized at 100 kb resolution. Orange boxes: A-A interactions in corresponding regions of Pro-E and Ortho-E; blue boxes: B-B interactions in corresponding regions of Pro-E and Ortho-E. **d**, Representative immunofluorescence images of H3K9me3 distribution in Pro-E and Ortho-E cells. **e**, Interaction length distribution of H3K9me3-HiChIP in Pro-E and Ortho-E. **f**, Respective percentage for intra- and inter-compartment interactions of H3K9me3-HiChIP in Pro-E and Ortho-E cells. **g**, (Top) PC1 value of Hi-C analysis indicating chromatin compartmentalization at chromosome 2 (0–60 Mb). Red: compartment A; blue: compartment B. (Middle) Corresponding H3K9me3-HiChIP interactions in Pro-E and Ortho-E cells. (Bottom) Corresponding H3K9me3-HiChIP contact matrices in Pro-E and Ortho-E cells. Juicebox parameters were balanced normalized at 100 kb resolution.

Since the canonical heterochromatin marker H3K9me3 is locolized within compartment B, we next performed immunofluorescence staining of H3K9me3 in Pro-E and Ortho-E cells. H3K9me3 localizes preferentially underneath the nuclear membrane with a “ring” like distribution in Ortho-E, while in Pro-E, H3K9me3 has a smear distribution throughout the nucleus (Fig. 2d and Extended Data Fig. 3a,b). However, surprisingly, CUT&RUN analysis of H3K9me3 revealed that the genomic localization of H3K9me3 was almost identical between Pro-E and Ortho-E stages (Extended Data Fig. 3c). Furthermore, the protein level of either H3K9me3 or H3K9me2 did not change significantly from Pro-E to Ortho-E stage (Extended Data Fig. 3d). These results demonstrate that the abundance of H3K9me3 marks do not change in terminal erythropoiesis, but genomic location distributes from entire nucleus to nuclear periphery, suggesting H3K9me3-marked heterochromatin may contribute to terminal erythroid chromatin compaction through three-dimensional reorganization.

We then performed *in situ* Hi-C followed by chromatin immunoprecipitation (HiChIP) of H3K9me3 to dissect the 3D heterochromatin organization^33^. We found a marked shift of H3K9me3-associated middle to long range loop length distribution during the transition from Pro-E to Ortho-E stage (Supplementary Table 2). Compared to the Pro- E stage, Ortho-E stage cells acquire more long-range interactions (>1Mb) but lose the middle-range interactions (between 0.1Mb and 1Mb) (Fig. 2e). The frequency of long interactions (>1Mb) increased from 38.13% to 54.04% in Ortho-E stage (Fig. 2e). Additionally, in consistent with increased interaction frequency among B compartments, Ortho-E cells show more inter-compartment interactions associated with H3K9me3 (Fig. 2f). For example, at the 0-60Mb region of chromosome 2, H3K9me3 associated compartments (compartment B) exhibit more inter-compartment interactions. The corresponding heatmap of H3K9me3-HiChIP also shows more long-range inter-compartment interactions (Fig. 2g).

Collectively, our results reveal that the heterochromatin undergo a conspicuous three-dimensional reorganization during terminal erythropoiesis, without discernible changes in the abundance and chromatin occupancy of heterochromatin histone marks (H3K9me3 and H3K9me2). The interactome of H3K9me3-marked heterochromatin is reorganized to gather at a ring-like subnuclear localization in proximity to the nuclear membrane and enriched for newly-formed long-range interactions greater than 1 Mb. Considering the overall chromatin compaction revealed by the Hi-C analysis (Fig. 1c), these results indicate that the newly generated long-range heterochromatin interactions may play a major role to achieve the highly condensed chromatin state in human terminal erythroid development.

### A large number of domain boundaries are disrupted in terminal erythroid chromatin compaction

We assessed the chromatin structure at the TAD level by multiple TAD-calling algorithms^19, 34^ and uncovered that the number of TADs dramatically decreased during terminal erythroid differentiation. The disruption of TADs is found in both non-cell-cycle-specific and G0/G1-sorted samples (Extended Data Fig. 4a). It is consistent with the larger average TAD size in Ortho-E, which indicates that small TADs are fused into larger ones (Extended Data Fig. 4b).

We also examined the strength of TAD boundaries by the insulation score. In consistent with decreased TAD number, the average boundary insulation scores are elevated, reflecting weakening of domain boundaries from Pro-E to Ortho-E (Fig. 3a). The average interaction heatmap at TAD boundaries showed higher frequency of cross-boundary interactions in Ortho-Es than in Pro-Es (Fig. 3b), and the average interaction heatmap at TAD regions showed decreased intra-domain interactions at the Ortho-E stage compared to those at the Pro-E stage (Extended Data Fig. 4c).

**Fig. 3:**
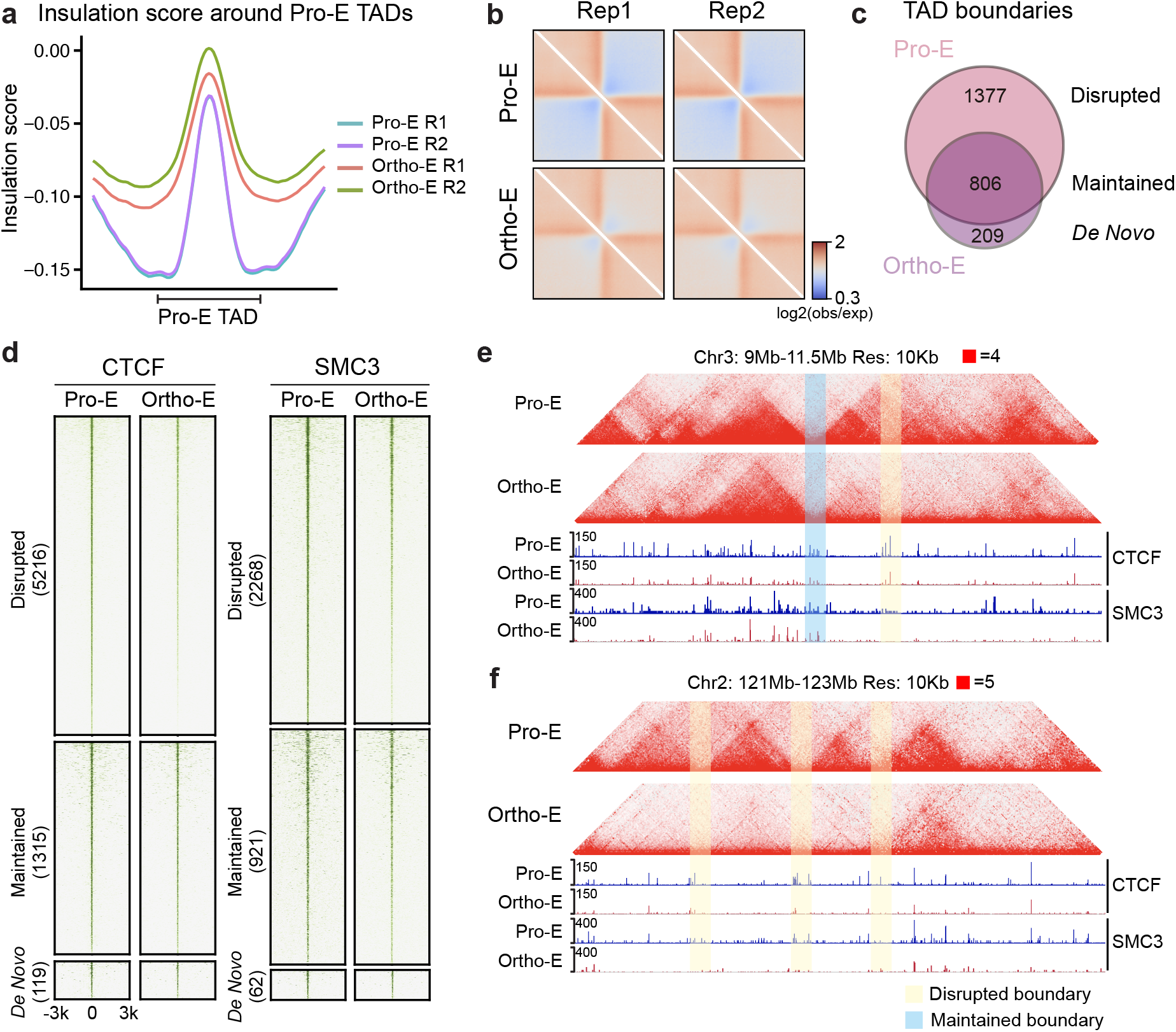
A vast number of chromatin domain boundaries are disrupted in terminal erythropoiesis. **a**, The average insulation scores at TADs and the neighboring regions (±0.5 TAD length) of Pro-E (cyan, purple) and Ortho-E (red, green). The analysis is based on TADs defined in the Pro-E stage. **b**, Heatmaps showing the normalized average interaction frequencies for all TAD boundaries (defined in Pro-E) and their neighboring regions (±0.5 TAD length) in Pro-E and Ortho-E. **c**, Venn plot showing the number of unique and overlapped TAD boundaries in Pro-E and Ortho-E. (Disrupted: the boundaries that are only present in Pro-E; maintained: the boundaries that are present in both Pro-E and Ortho-E; de novo: the boundaries that are only present in Ortho-E.) **d**, Density maps of CTCF and SMC3 signals at the disrupted, maintained and de novo formed TAD boundaries in the Pro-E and Ortho-E stage by CUT&RUN assay. **e**, (Top) Hi-C contact matrices for chromosome 3 (9–11.5 Mb) of Pro-E and Ortho-E cells; (bottom) CTCF and SMC3 binding at the corresponding region. (Blue shadow: maintained boundary region; yellow shadow: disrupted boundary region) Juicebox parameters were balanced normalized at 100 kb resolution. **f**, (Top) Hi-C contact matrices for chromosome 2 (121–123 Mb) of Pro-E and Ortho-E cells; (bottom) CTCF and SMC3 binding at the corresponding region. (Blue shadow: maintained boundary region; yellow shadow: disrupted boundary region) Juicebox parameters were balanced normalized at 100 kb resolution.

We next classified the TAD boundaries present in Pro-Es into three categories based on their status of maintenance in Ortho-Es by using the cword method at 20kb resolution: “maintained boundaries” (806, 34%) are stable during terminal erythroid stages; “disrupted boundaries” (1377, 58%) and “*de novo* formed boundaries” (209, 8%) represent those disappeared and newly-formed boundaries during terminal erythropoiesis, respectively (Fig. 3c). The average insulation scores of disrupted, maintained and *de novo* formed boundaries increased, unchanged and decreased, respectively (Extended Data Fig. 4d).

To understand the molecular basis underlying the three categories of TAD boundaries, we examined their corresponding CTCF and cohesin binding by CUT&RUN assay, as CTCF and cohesin are asscioated with domain boundary insulation^35, 36^. The overall binding events of CTCF and SMC3 declined in terminal erythropoiesis (Extended Data Fig. 4e,f). Nonetheless, it is noteworthy that CTCF and SMC3 binding on “disrupted” boundaries decreased markedly, but remained relatively stable on “maintained” and “*de novo* formed” boundaries (Fig. 3d-f). These results show that the weakened boundaries in Ortho-Es are associated with CTCF and SMC3 binding loss.

Erythroid terminal differentiation involves a massive down-regulation of proteome^37^, and the protein level of CTCF and cohesin subunits have been shown to regulate boundary strength and chromatin architecture in cell lines^35, 38^. We then examined the protein level of CTCF and cohesin subunits (SMC3, RAD21) in terminal erythroid stages. Interestingly, protein expression of SMC3 and RAD21 were constant, whilst CTCF showed a ∼40% decrease in Poly-E and Ortho-E (Extended Data Fig. 4g). Since the chromatin remodeling activity of ISWI complex and CHD families can regulate the deposition of CTCF by moving nucleosomes^39, 40^, we next detected the expression of SNF2H, BRG1 and CHD4, which are the ATPase subunit of ISWI, SWI/SNF and CHD4 chromatin remodeling complexes, as well as the cellular ATP content in terminal erythroid stages. We found that the level of SNF2H, BRG1 and CHD4 greatly declined, so did the cellular ATP content (Extended Data Fig. 4g,h). Furthermore, our MNase-seq also showed elevated nucleosomal signal at the regions where CTCF binding dropped (Extended Data Fig. 4i). These results suggest that loss of CTCF and SMC3 binding at the disrupted boundaries might be associated with a series of cellular events during terminal erythropoiesis including down-regulation of transcription/translation activity and restricted energy state.

Together, we have observed a striking decrease of TAD boundary strength during terminal erythroid development. Most TAD boundaries are disrupted, while select TAD boundaries are maintained in terminal erythropoiesis. The boundary deficiency of Ortho-E is associated with loss of CTCF and SMC3 binding, likely resulting from restricted cellular energy.

### Select domain maintenance in the Ortho-E stage is associated with transcription competent state and erythroid gene expression

Since a select group of TAD boundaries remain intact, while a large number of TADs are disintegrated in terminal erythroid stages, we reason that the maintained TADs may exhibit distinct molecular features from the disrupted ones. Firstly, we assessed the strength of TADs using “TAD score” (domain score) which is the ratio of intra-TAD interactions to total interactions associated with a defined TAD region (Intra- plus Inter- TAD interactions) (Fig. 4a)^41^. We next calculated the overall TAD score of Pro-E and Ortho-E stage based on TADs defined in the Pro-E stage, and found that the average TAD score decreased in Ortho-E cells, meaning the global TAD structures were attenuated in terminal erythropoiesis (Fig. 4b). To dissect the molecular basis underlying the differential regulation of Pro-E TADs in Ortho-E stages, we classified the Ortho-E TADs into 10 groups, based on the TAD score ratio of Pro-E to Ortho-E stage: “Stable TADs” (Group 1&2) indicate the ones that have increased or maintained TAD scores during terminal stages; “Disrupted TADs” (Group 9&10) represent the ones with fiercely decreased TAD scores during terminal stages (Fig. 4c and Extended Data Fig. 5a).

**Fig. 4:**
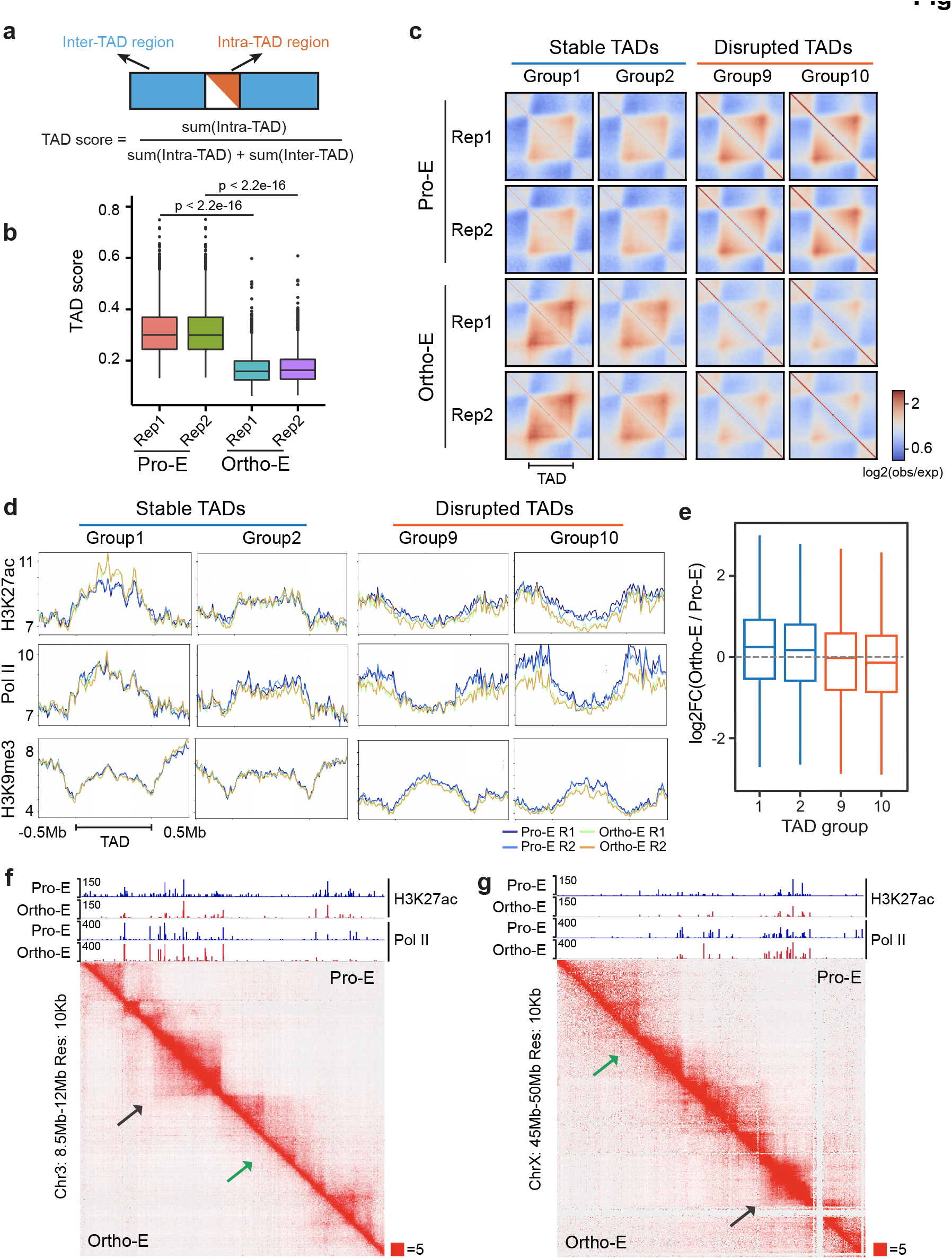
Domain maintenance in the Ortho-E stage is determined by chromatin activity and erythroid gene expression. **a**, Schematic description for the calculation of TAD score (domain score). **b**, TAD score distribution of Pro-E and Ortho-E, based on TAD regions defined in Pro-E. **c**, Heatmaps showing the normalized average interaction frequencies for TADs and the neighboring ±0.5TAD regions. The TADs are grouped by the TAD score ratio (TAD score of Ortho-E to that of Pro-E). Group1&2 represent the most “Stable” TADs; group 9&10 represent the most “Disrupted” TADs. **d**, Distribution of H3K27ac, Pol II and H3K9me3 signal by CUT&RUN assay in TADs and the neighboring ±0.5 Mb regions. The TADs are grouped by their TAD score ratio. Group1&2 represent the most “Stable” TADs; group 9&10 represent the most “Disrupted” TADs. **e**, Fold change (FC) of gene expression within the Stable (Group1&2) vs. Disrupted (Group9&10) TADs. **f**, Representative Hi-C interaction matrices and the corresponding H3K27ac and Pol II signals by CUT&RUN assay at the “Stable”(black arrow) and “Disrupted”(green arrow) TAD regions in Chr3: 8.5Mb-12Mb. Juicebox parameters were balanced normalized at 10 Kb resolution. **g**, Representative Hi-C interaction matrices and the corresponding H3K27ac and Pol II signals by CUT&RUN assay at the “Stable” (black arrow) and “Disrupted” (green arrow) TAD regions in ChrX: 45Mb-50Mb. Juicebox parameters were balanced normalized at 10 Kb resolution.

We then analyzed the enrichment of active histone markers — H3K27ac in stable vs. disrupted TADs of different groups. Our results showed that H3K27ac are preferentially enriched in the “Stable TADs”, with Group 1 having the most pronounced signals. Furthermore, H3K27ac signals are greater in the Group 1 TADs of Ortho-Es compared to those of Pro-Es (Fig. 4d and Fig. 4f,g). Consistent with the deposition of H3K27ac, RNA Pol II exhibits preferential occupancy in the stable TADs, with Group 1 and 10 having the highest and lowest Pol II occupancy, respectively. (Fig. 4d and Fig. 4f,g). On the contrary, H3K9me3 marks are not differentially distributed among distinct TAD groups (Fig. 4d). As CTCF and cohesin are involved in the formation of intra-TAD structures and chromatin loops^42, 43^, we next investigated whether the presence of CTCF and SMC3 is indicative of the stability of TADs. The results showed that CTCF and SMC3 are highly enriched in the “Stable TADs”, especially Group 1&2, but showed an invert occupancy pattern in the “Disrupted TADs”. It is also noted that in the “Stable TADs”, SMC3 of Pro-Es has a higher enrichment at intra-TAD regions than that of Ortho-Es, whilst in the “Disrupted TADs”, SMC3 of Pro-Es has a higher enrichment at the boundary regions than that of Ortho-Es. On the other hand, CTCF shows stable deposition when comparing the signals of Pro-Es and Ortho-Es in different TAD groups, except slightly decreased signal at the disrupted boundary regions in Ortho-E (Extended Data Fig. 5b).

Having established that active chromatin marks and Pol II are associated with TAD maintenance in the Ortho-E stage, we next examined whether transcription activity serves as a key determinant to the maintenance of TADs by analyzing the changes of gene expression within different TAD groups. Gene activation in the Ortho-E stage was highly correlated with TAD maintenance (Group 1&2); similarly, gene repression was associated with TAD disruption in Ortho-E cells (Group 9&10) (Fig. 4e). Gene Ontology analysis shows the up-regulated genes in the “Stable TADs” represent biological processes such as autophagy (Supplementary Tables 3-4), while repressed genes in the “Disrupted TADs” are associated with various RNA-related pathways (Extended Data Fig. 5c and Supplementary Tables 5,6).

Taken together, we have uncovered that TAD structures are differentially disrupted, with only a select group of TADs being maintained throughout terminal erythropoiesis. We dissected the molecular features of the “Stable TADs” in the Ortho-E stage and found that the stable TADs are enriched for H3K27ac, Pol II, cohesin and CTCF binding. Furthermore, our analysis showed that the up-regulated genes within the “Stable TADs” are enriched for the autophagy pathway, which is critical for terminal erythropoiesis. These data indicate the signature of active chromatin state and transcription competence serve as the key determinant for the differential maintenance of chromatin domains in terminal erythroid development, and the selectively maintained domains may play more important roles in terminal erythroid functions. These findings may also imply possible involvement of erythroid transcription regulators in the maintenance of TADs at terminal development.

### GATA1 regulates the chromatin architecture maintenance in terminal erythropoiesis

As the select maintenance of TADs is associated with transcriptionally active chromatin state, we reason that transcription regulators may also contribute to the regulation of chromatin architecture. We conducted the Assay for Transposase-Accessible Chromatin with high-throughout sequencing (ATAC-seq) and *de novo* motif analysis, which unveiled that the motifs of two erythroid master transcription factors, GATA1 and KLF1, remain relatively accessible during the Pro-E to Ortho-E transition (Extended Data Fig. 6a), but the accessibility of GATA1 motifs diminished in the late Ortho-E stage^26^. GATA1 is a master transcription factor of erythroid development^44^. Loss of GATA1 in the terminal erythroid stages results in abnormal compression of the nucleus, which in turn affects enucleation^45^.

We next investigated GATA1 and KLF1 binding in different groups of TADs by CUT&RUN assay. Similar to the deposition of H3K27ac and Pol II, occupancy of GATA1 and KLF1 is highly enriched in the “Stable TADs”, especially Group 1, and these two factors have an inverse correlation of occupancy in the “Disrupted TADs” (Group 9&10) (Fig. 5a). To understand whether GATA1 and KLF1 mediate maintenance of chromatin architecture via promoting chromatin looping, we performed GATA1 and KLF1 HiChIP in terminally differentiated erythroblasts of Day 12 in the *in vitro* erythroid culture system. Similar to the occupancy in the CUT&RUN assay, the anchor points of GATA1 and KLF1 by HiChIP are prodominantly located in the “Stable TADs”, and the anchor point signals are negatively correlated with the genomic regions of the disrupted TADs (Fig. 5b-e and Extended Data Fig. 6b). These results indicate that master transcription factors GATA1 and KLF1 are likely involved in maintaining the functionally important chromatin domains during terminal erythropoiesis.

**Fig. 5:**
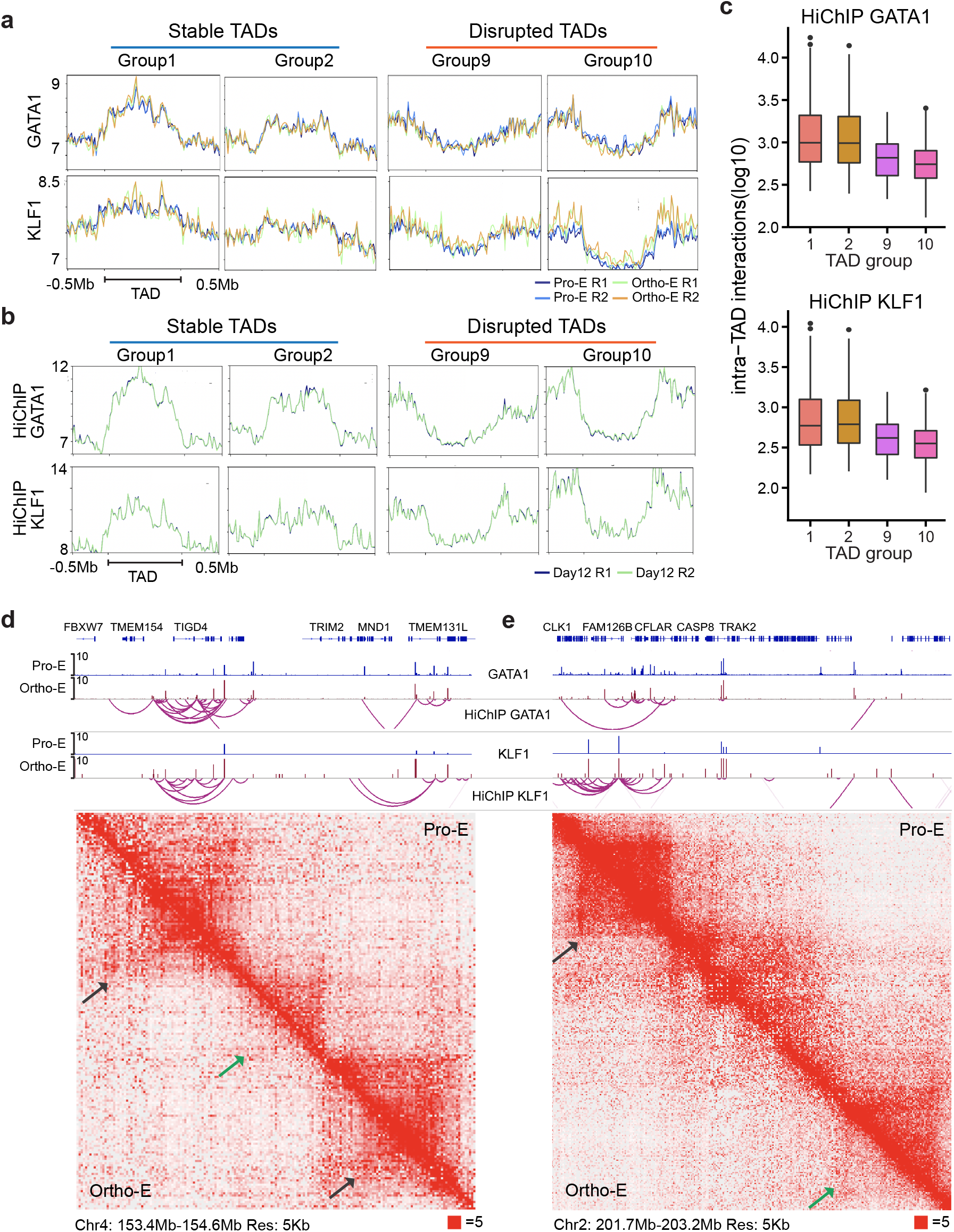
Erythroid master regulators GATA1 and KLF1 are strongly associated with maintained chromatin domains. **a**, GATA1 and KLF1 signal distribution (CUT&RUN) in TADs and the neighboring ±0.5 Mb regions. The TADs are grouped by their TAD score ratio. Group 1&2 represent the most “Stable” TADs; group 9&10 represent the most “Disrupted” TADs. **b**, Signal distribution of GATA1 and KLF1 at the loop anchors of HiChIP assay in TADs and the neighboring ±0.5 Mb regions. The TADs are grouped by their TAD score ratio. Group 1&2 represent the most “Stable” TADs; group 9&10 represent the most “Disrupted” TADs. **c**, (Top) GATA1 or (bottom) KLF1 mediated intra-TAD interaction frequency within the Stable (Group 1&2) vs. Disrupted (Group 9&10) TADs. **d**, Representative Hi-C interaction matrices and the corresponding GATA1 and KLF1 signals, along with chromatin loops mediated via these two factors by HiChIP assay at the “Stable”(black arrow) and “Disrupted”(green arrow) TAD regions in Chr4: 153.4 Mb-154.6 Mb. Juicebox parameters were balanced normalized at 5 Kb resolution. **e**, Representative Hi-C interaction matrices and the corresponding GATA1 and KLF1 signals, along with chromatin loops mediated via these two factors by HiChIP assay at the “Stable”(black arrow) and “Disrupted”(green arrow) TAD regions in Chr2: 201.7 Mb-203.2 Mb. Juicebox parameters were balanced normalized at 5 Kb resolution.

To test the function of GATA1 and KLF1 in maintaining terminal erythroid chromatin architecture, we knocked out GATA1 and KLF1 via CRISPR/Cas9 in sorted primary Pro-E cells (Fig. 6a and Extended Data Fig. 7a-c). Knocking out GATA1 abrogated the expression of CD235a and severely affected differentiation of Pro-E, while knocking out KLF1 did not affect the expression of CD71 and CD235a, resulting in a minor cell morphological phenotype (Fig. 6b and Extended Data Fig. 7d). We performed Hi-C after knocking out GATA1 and KLF1 for 72 hours (h), when the control cells are mostly differentiaed into the Ortho-E stage. The overall TAD score and the global intra-TAD interaction diminished after abrogation of GATA1 (Fig. 6c). Consistently, the number of TADs decreased after knocking out GATA1 (Fig. 6d). For example, at genomic regions such as Chr2: 60.5Mb-63Mb and Chr3: 182Mb-184Mb, knocking out GATA1 impaired maintenance of intra-TAD interactions in the “Stable TADs” (Fig. 6e,f and Extended Data Fig. 7e,f). However, we did not observe notable changes of TADs or intra-TAD interactions after ablation of KLF1 (Extended Data Fig. 7g,h). Also, we did not observe obvious changes of TAD structure at 48h post knocking-out of GATA1 (data not shown). These data indicate that the function of GATA1 in maintaining TAD structures is more prominent in the late stage (Ortho-E) of erythroid development, during which time TAD structures undergo global attenuation.

**Fig. 6:**
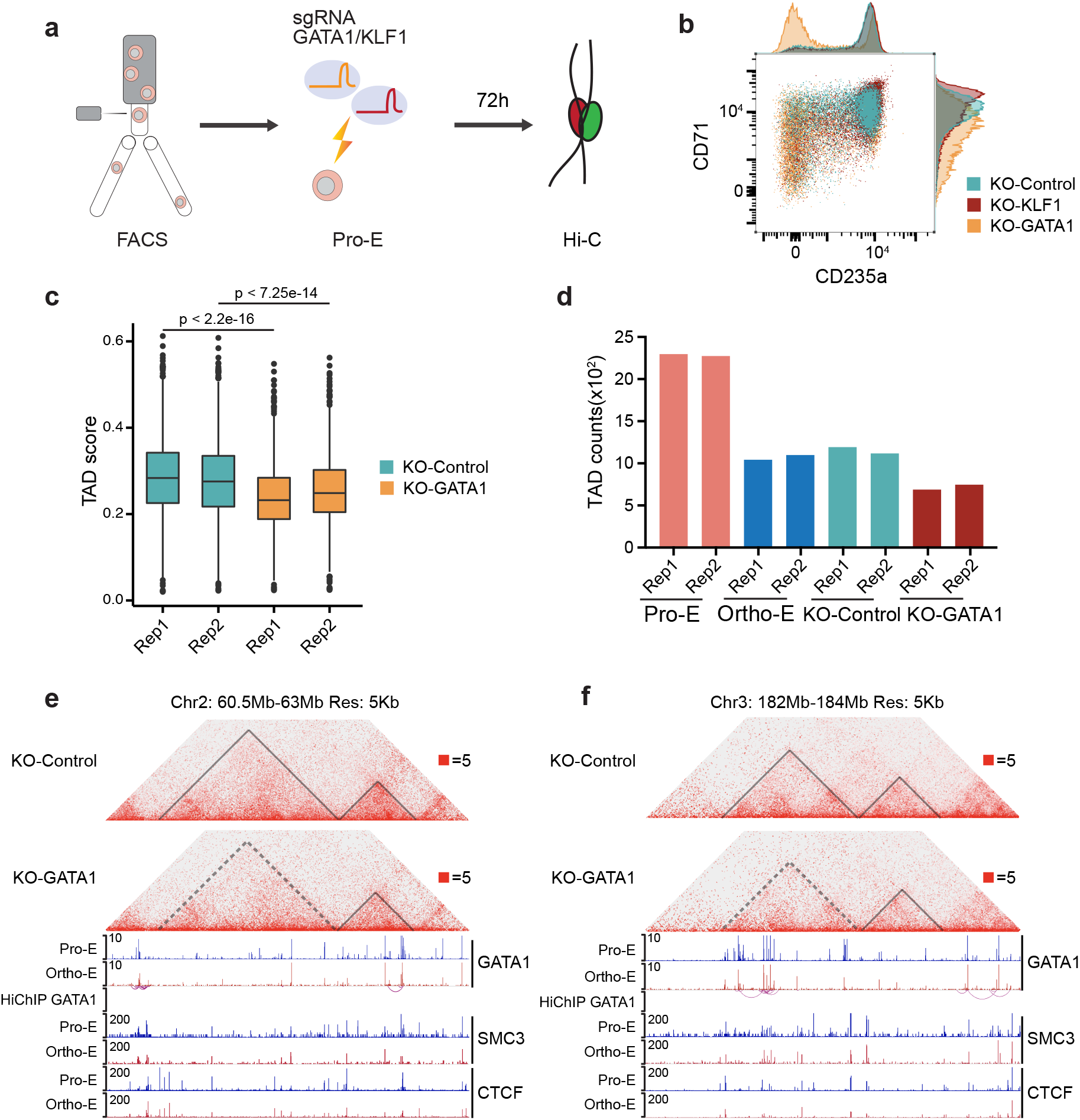
GATA1 is essential in safeguarding select chromatin domains in terminal erythropoiesis. **a**, Experimental scheme for CRISPR/Cas9 knockout of GATA1 or KLF1 in human primary Pro-E cells. **b**, Erythroid differentiation monitored by the expression of CD71 and CD235a after knocking out by GATA1, KLF1 or control sgRNA for 72 h. **c**, TAD score distribution of all TADs after knocking out by GATA1 or control sgRNA. TAD regions are defined in the Pro-E stage. **d**, The number of TADs in Pro-E, Ortho-E and Pro-E cells knocking out GATA1 or control sgRNA for 72 h. **e**, (Top) Hi-C contact matrices of chromosome 2 (60.5–63 Mb) after control or GATA1 knockout; (middle) the corresponding GATA1 CUT&RUN signal and HiChIP loops in Pro-E and Ortho-E; (bottom) the corresponding SMC3 and CTCF CUT&RUN signals in Pro-E and Ortho-E. Juicebox parameters were balanced normalized at 5 Kb resolution. Dashed triangle shows that the disrupted TAD resulting from knocking-out of GATA1. **f**, (Top) Hi-C contact matrices of chromosome 3 (182–184 Mb) after control or GATA1 knockout; (middle) the corresponding GATA1 CUT&RUN signal and HiChIP loops in Pro-E and Ortho-E; (bottom) the corresponding SMC3 and CTCF CUT&RUN signals in Pro-E and Ortho-E. Juicebox parameters were balanced normalized at 5 Kb resolution. Dashed triangle shows that the disrupted TAD resulting from knocking-out of GATA1.

We next delved into the potential mechanisms of GATA1 safeguarding select TADs in terminal erythroid stages. Unlike CTCF and YY1, which directly mediate chromatin looping by forming homodimers, GATA1 and KLF1 were reported to mediate chromatin looping via recruiting a coregulator that mediates looping, e.g., LDB1^46, 47^. We analyzed the occupancy of GATA1, KLF1, CTCF and SMC3 in maintained vs. disrupted TADs, and found that GATA1 and KLF1 co-localized with CTCF and SMC3 in the maintained TADs (Fig. 6e,f and Extended Data Fig. 7i). Since knocking out KLF1 at the terminal erythroid stages did not yield notable phenotypes, our results suggest that GATA1 is the major regulator that mediates TAD maintenance in the Ortho-E stage in a CTCF and SMC3 dependent manner.

Our results showed that abrogation of GATA1 in terminal erythroblasts impairs the maintenance of select TADs in terminal erythroid development. To this end, we uncovered a previously unidentified function of GATA1 in safeguarding higher-order chromatin structure while there is a tremendous disruption of global TAD structures during chromatin compaction of terminal erythropoiesis.

## Discussion

The major difference of erythropoiesis in mammals and non-mammalian vertebrates is that mature red blood cells of vertebrates contain an intact nucleus, while mammalian erythroblasts expel their nuclei to produce enucleated mature red blood cells. During terminal erythroid development of both mammals and non-mammalian vertebrates, cell nuclei undergo significant compression^48, 49^. However, how the chromatin goes through multi-dimensional rearrangement to compress 70-80% of the nuclear volume while maintaining essential functions, such as transcription, remain elusive. In non-mammalian vertebrates, such as chicken, which have mature red blood cells containing an intact nucleus, the erythrocyte genome exhibits distinct A/B compartments and has no canonical TADs^50^.

Here in this study, for the first time, we uncovered that *de novo* ultra-long-range interactions established by heterochromatin reorganization play a major role in human erythroid chromatin compression. Meanwhile, more than 63% of TAD boundaries are drastically disrupted, which is associated with loss of CTCF and SMC3 binding in the terminal development of human erythropoiesis. Interestingly, we also identified a group of TADs that are selectively maintained in the Ortho-E stage, which represents the last developmental stage before enucleation. In contrast to the mature chicken erythrocytes genome that lacks canonical TAD structures, about 37% of TAD structures remain intact till the end of human terminal erythropoiesis.

By systematically examining the molecular characteristics of the maintained TADs, our results reveal that these maintained TADs are enriched for histone marks and factors indicative of active chromatin state, including H3K27ac and Pol II. Furthermore, we have identified a previously unappreciated role of the master transcription factor, GATA1, in safeguarding the chromatin domains that are associated with terminal erythroid functions, such as autophagy, while occupancy of GATA1 is inversely associated with the disrupted chromatin regions. Our functional analysis demonstrated that knocking out GATA1, but not KLF1, in the Pro-E stage abrogated terminal erythroid chromatin architecture.

Terminal erythroblasts have a shorter cell cycle than normal cells, and the distinct properties of terminal erythroid cell cycle were reported before^30^. Chromosome folding is regulated periodically as cells progress through the cell cycle, and dynamics of chromatin architecture have been well characterized in cell cycle transition. During prometa to G1 phase transition, rapid establishment of A/B compartments was observed, followed by their gradual intensification and expansion. Contact domains form following the “bottom up” principle— that the small domains fuse into a large TAD domain^32, 51, 52^. In the current study, we isolated G0/G1 cells of human terminal erythroblasts to avoid the effects of cell cycle on Hi-C assay, even though our Hi-C analyses comparing cell cycle specific vs. non-specific samples revealed consistent results. Compared with the prometa phase of other normal cell types which is characterized by blurry TAD and compartment structures, human terminal erythroblasts exihibit distinct TAD structures that undergo immense disruption, leaving a select number of maintained TADs until the end of erythroid differentiation; moreover, A/B compartment partition also remains constant throughout terminal erythropoiesis (Extended Data Fig.2, 3). Collectively, these findings demonstrate that the 3D genome structure of terminally differentiated erythroblasts is largely distinct from that of mitotic cells. Nonetheless, there are questions which remain to be answered. Chromatin compaction happens progressively in terminal erythroid development, but how does the chromatin compaction accumulate and propagate through the last 3-5 cell cycles before cell cycle exit and mammalian erythroid enucleation? We propose a model that the final chromatin compaction state of Ortho-E is an accumulative result of partial decompaction after each of the last 3-5 cycles. It is likely due to the decrease of cellular energy (ATP concentration) at terminal erythroid stages, ATP-dependent cellular activities, such as chromatin remodeling, is substantially affected, leading to incomplete chromatin decompaction and accumulated chromatin compaction after each terminal cell cycle (Extended Data Fig. 4g,h). This is reminiscent of B cell activation, in which global histone acetylation and ATP synthesis lead to genome decompaction^53^. However, even though HDAC proteins were reported to increase in terminal erythropoiesis^9^, no global declination of histone acetylation in terminal erythropoiesis was observed^54^, suggesting that chromatin compaction and the corresponding architecture changes are likely controlled by additional intricate mechanisms (Extended Data Fig. 8).

Recent studies have identified a plethora of factors that can affect higher-order chromatin structure, such as Pol II, and certain transcription factors, such as YY1 and MyoD^42, 55^, which can form homodimers to mediate long-range interactions among chromatin regions^55^. Erythroid master transcription factor GATA1 has been reported to mediate chromatin interactions through looping factors (such as LDB1)^46, 47, 56^. In addition, previous research revealed that certain GATA1 binding sites localized at key hemapotoietic genes can exert a bookmarking function during cell cycle progression^27^. Our results demonstrated that GATA1 occupancy is highly enriched in the maintained TADs of Ortho-Es, which are also co-localized by CTCF and SMC3. Moreover, our functional assay knocking out GATA1 showed a significant disruption of TAD structures in primary terminal erythroblasts. Given that GATA1 has been shown to recruit chromatin remodeling complexes and mediate chromatin looping via LDB1-containing complexes and CTCF, we reason that GATA1 might play a critical role in allocating limited cellular energy for essential chromatin remodeling activities as well as CTCF and SMC3 binding to safeguard select chromatin domains in highly-compacted terminal erythroid nuclei, maintaining required nuclear activities, including transcription (Extended Data Fig. 8). To this end, we elucidated the molecular fundamentals of a development-driven chromatin condensation process using primary human terminal erythropoiesis system, and uncovered that transcription competence serves as a key determinant in managing higher-order nuclear architecture to maintain accessibility of select essential domains for developmental functions.

## Methods

### *Ex vivo* CD34+ cells erythroid culture

Cord blood cells were obtained from the Cord Blood Bank of Beijing. CD34+ cells were purified by using MACS MicroBead kit (Miltenyi, Cat. # 130-100-453) and cultured by a two-phase erythroid differentiation method modified from a previously published method^57^. The base medium contains IMDM, 10% FBS (Gibco, Cat. #10099141), 300 μg/ml holo-transferrin (Sigma, Cat. #T0665), 5% human AB serum (Wokavi Biotech, Beijing), 10 ng/ml heparin (Sigma, Cat. #H3149), 10 μg/ml insulin (Sigma, Cat. #I9278), 2 mM L-Glutamine (Gibco, Cat. #25030081) and 3 IU/ml erythropoietin (Amgen, Cat. #55513-144-10). Phase-I medium was supplemented with 50 ng/ml hSCF (StemCell, Cat. #78062) and 10 ng/ml hIL-3 (StemCell, Cat. #78042). Phase-II medium was supplemented with 50 ng/ml hSCF only. Cells were maintained in Phase-I until day 8 and then transferred to Phase-II medium for the rest of the culture period.

### Isolation of primary human erythroblasts at distinct terminal differentiation stages

Terminally differentiated erythroid cells were sorted by fluorescence-activated cell sorting (FACS), based on a published method modified for our purpose^29^. Briefly, at day 12 of CD34+ cells erythroid culture, cells were spun down and resuspended in the staining buffer (2% FBS in PBS) and dead cells were removed by Annexin-V MicroBeads(Miltenyi, Cat.#130-090-201); then the following antibodies were added as per 1,000,000 cells: 5μl anti-CD235a (APC, eBioscience, Cat.# 17-9987-42), 5μl anti-CD71 (FITC, eBioscience, Cat.#11-0719-42), 6μl anti-CD45 (PerCP-Cy5.5, BD, Cat.#564105), 5μl anti-CD36 (APC-Cy7, BioLegend, Cat.#336214), 5μl anti-CD117 (PE, eBioscience, Cat.#12-1178-42), 5μl anti-CD105 (PE-CF594, BD, Cat.#562380). After staining, cells were washed with the staining buffer and 2 μg/ml Hoechst33342 was added. FACS sorting was performed on BD FACSAria^TM^ II SORP. After gating G0/G1 cells by Hoechst33342 staining, cell populations of distinct differentiation stages were isolated by the following cell surface markers: Pro-E cells are CD235a^+^CD71^+^CD45^low^CD36^+^CD117^high^CD105^high^; Baso-E cells are CD235a^+^CD71^+^CD45^low^CD36^+^CD117^dim^CD105^dim^; Poly-E cells are CD235a^+^CD71^+^CD45^low^CD36^+^CD117^-^CD105^-^FSC^high^; Ortho-E cells are CD235a^+^CD71^+^CD45^low^CD36^+^CD117^-^CD105^-^FSC^low^. The analytical flow cytometry was performed on BD Fortessa SORP by the same staining method.

### CRISPR–Cas9 mediated gene knockout in human primary Pro-E cells

Cas9–sgRNA RNP complexes were assembled as follows. For each knockout experiment, 300 pmol of Cas9 protein (IDT) and 500 pmol of modified sgRNA (Genescript) were mixed and incubated for 20 min at room temperature. One million sorted Pro-E cells were resuspended with 100 μl solution P3 for primary cells (Lonza); preassembled RNP were then added to the cell suspension for nucleofection. Nucleofection was performed in a 4D-Nucleofector X unit (Lonza) using program EO-100. Cells were centrifuged and transferred to 2 ml Phase-II medium for continuing culture under erythroid differentiation conditions. To assess genome editing efficiency, cells were lysed for genomic DNA extraction. Primers spanning the edited sites were used to amplify the genomic region. A pair of gRNAs or three gRNAs were introduced to perform knocking-out of KLF1 and GATA1, respectively. The sequences of gRNAs and genotyping primers are listed in the Supplementary Table 7^45, 58^.

### *In situ* high-through chromosome conformation capture (Hi-C) assay

Hi-C assay was carried out using the in-situ method as described before with minor modifications^17^. Half a million FACS-purified primary erythroblasts were used for each reaction. In brief, after fixation with 1% formaldehyde, cells were digested overnight at 37°C using 50U of MboI (NEB, Cat. #R0147). After biotin filling, proximity ligation was carried out overnight at 25 °C with 1600U T4 DNA Ligase (Vazyme, Cat. # N103-01). After reverse-crosslinking, DNA was purified and sheared to 300-500bp fragments using Covaris M220 sonicator. Ligation fragments containing biotin were immobilized on MyOne Streptavidin C1 beads (ThermoFisher Cat. #: 65602), and then library preparation was performed using the DNA library preparation kit (Vazyme, Cat.# ND607) as per the manufacturer’s protocol. DNA was then subjected to double-size selection using DNA Clean Beads (Vazyme, Cat. # N411) in order to isolate fragments between 300 and 600bp. Two biological replicates per condition were sequenced in a HiSeq X-Ten (PE150, Illumina) or NovaSeq 6000 (S4, PE150, Illumina) by Novogene (Beijing).

### *In situ* Hi-C data analysis

Hi-C data was processed and analyzed as previously published^17^. The paired-end raw reads of Hi-C libraries were generated, and HiC-Pro (version 2.10.0) was used to map, filter invalid reads and process ICE normalization with default parameters^59^. Human genome hg19 was used as reference. Then valid read pairs were binned at a series of different resolutions. The “sample.hic” file was created by using the HiC-Pro additional scripts and visualized through the Juicebox (version 1.9.0) software^60^. The DLR (distal to local) and ICF (inter-chromosome frequency) were calculated using Homer^61^. Besides, the contact probability as a function of distance was calculated by using pairsqc scripts (https://github.com/4dn-dcic/pairsqc), which is a tool for quality control of Hi-C data. Research in the past provided^62^ that P(s)∼s^-1^ and P(s)∼s^-0.5^ represent the fractal globule and mitotic states.

### Analyses of TADs and TAD boundaries

The 20kb resolution matrix was used to performed TAD calling. We called TADs by using three different methods: insulation score^63^, Homer^61^ and Arrowhead^60^. The default parameters were applied in the three methods. It is noteworthy that analyzing the same samples using different methods of distinct algorithms could result in varied TAD counts, but the main conclusions remain valid despite the bioinformatic methods used. To identify TAD boundaries and TADs, we used the script “matrix2insulation.pl” with the default parameters^63^. The strength of TAD boundaries was derived from the results of insulation score. We merged the boundaries found in two replicates of the same cell stage, i.e. Pro-E or Ortho-E, and then compared the boundaries of Pro-E and those of Ortho-E to identify the developmental stage specific boundaries. To compare the insulation score of boundaries, we used the Pro-E TADs as baseline of comparison and extended our selection to be a 2Mb region covering and surrounding the TADs, and plotted the averaged insulation score in Pro-E and Ortho-E samples. The aggregated boundary or aggregated TAD analyses were performed for different groups of TADs using coolpup.py ^64^. Otherwise, the TAD score was defined using the ratio of “sum of intra-TAD interaction” to “sum of inter-TAD and intra-TAD interactions”. Then TAD scores ratio (Pro-E/Ortho-E) were ranked to select significantly different TAD regions between Pro-E and Ortho-E. Deeptools was used to generate enrichment profiles of transcription factors and histone modifications in different TAD groups^65^.

### Identification of chromatin compartments and analysis

As in previous studies, principal component value was used to call compartment A/B by using KR normalization with the 100kb and 500kb resolution for each sample using Juicertools ^60^. The chromatin can be assigned as compartment A or compartment B according to the positive or negative PC1 values, respectively. We validated the compartment assignment by assessing the level of gene expression and histone modifications in compartment A/B. The WashU Epigenome Browser (http://epigenomegateway.wustl.edu/) and IGV (Integrative Genomics Viewer) were used for the visualization of compartments ^66, 67^. The dynamic compartment switching from Pro-E to Ortho-E was calculated based on the compartments that were identified in both replicates. If a compartment was switched from A to B during the transition of Pro-E to Ortho-E, then the compartment was designated as “AB” and vice versa. Otherwise, “AA” or “BB” were used to indicate the stable compartments during erythropoiesis. The PC1 value was ranked and the top 20% A-A interactions and top 20% B-B interactions were used for quantification.

### Cleavage Under Targets and Release Using Nuclease (CUT&RUN) assay

CUT&RUN experiments were carried out as described with modifications^68^. Briefly, 100,000 cells were captured with BioMagPlus Concanavalin A (Bang Cat. # BP531) for 20 min at room temperature. Permeabilization of cells and binding of primary antibodies were performed for 2 hours at 4°C. After washing away unbound antibody with the washing buffer, protein A-MNase was added at a 1:1000 ratio and incubated for 2 hours. Cells were washed again and placed in a 0°C dry bath metal block. Then CaCl_2_ was added to a final concentration of 2 mM to activate protein A-MNase. The reaction was carried out for 30 min and stopped by addition of equal volume of 2XSTOP buffer. The protein-DNA complex was released by centrifugation. DNA was extracted by NEB PCR Cleanup Kit (NEB Cat. # T1030), followed by quality control steps using Qubit fluorometer and bioanalyzer. Protein A-MNase was expressed and purified from BL21(DE) carrying pET-pA-MN (Addgene: 86973).^68–70^

The CUT&RUN library was prepared using the DNA library preparation kit (Vazyme, Cat. # ND607) following the manufacturer’s protocol with minor modifications^71^. Briefly, the temperature of dA-tailing was decreased to 50°C to avoid DNA melting, and the reaction time was increased to 1 hour. After adaptor ligation, 2X volume of DNA Clean beads (Vazyme, Cat. # N411) was added to the reaction to ensure high recovery efficiency of short fragments. After 14 or 15 cycles of PCR amplification, the reaction was cleaned up with 1.2X volume of DNA Clean beads. The adaptor/PCR dimers were removed by E-Gel NGS Size Select II gel (Thermo Cat.# G661012). Two biological replicates per condition were sequenced in a HiSeq X-Ten (PE150, Illumina) by Novogene (Beijing).

### CUT&RUN data analysis

Raw sequence data of CUT&RUN were trimmed to the position of barcode using trim_galore (version 0.4.4_dev) and the reads were aligned to hg19 version of human genome with bowtie2 (version 2.3.4.1)^72^. Duplicated reads were removed through Picard tool (http://broadinstitute.github.io/picard/) and the low quality of reads (MAPQ < 30) were also removed. Reads with the cutoff of inset size < 120bp were selected for transcription factors occupancy and H3K27ac histone modification as previous studies. For other histone modifications, 150-500bp insert size reads were used for analysis^68, 73^. The bigwig files were generated by bamCoverage^65^ to visualize peaks in IGV^67^ or the WashU browser^66^. MACS2 with default parameters were used for peak calling^74^. At least two replicates of each CUT&RUN sample were used for analysis. The difference of transcription factors occupancy in Pro-E and Ortho-E was calculated using the diffbind R package.

### RNA purification and RNA-seq experiment

RNA was extracted with TRIzol and cleaned with RNeasy MinElute Cleanup Kit (Qiagen, Cat.# 74204). For RNA-seq, RNA samples were quantified and the quality was evaluated by Agilent 2100 Bioanalyzer. 1 μg of total RNA was used to prepare RNA-seq libraries with ribosomal RNA depletion using TruSeq Stranded Total RNA Library Prep Kit (Illumina, RS-122-2201). Two biological replicates per condition were sequenced in a HiSeq X-Ten (PE150, Illumina) by Novogene (Beijing).

### RNA-seq data analysis

RNA-seq data were subjected to quality control procedure by FastQC (version 0.11.7). Hisat2 (version 2.1.0)^75^ was used to align the paired-end raw data to hg19 version of human reference genome, and HTSeq was used to estimate the reads counts^76^. The commands were “hisat2 -x hg19 -t -p 8 -1 fq1 -2 fq2 --rna-strandness RF -S sample.sam, htseq-count -f bam -s reverse sample.bam annotation.gtf > output.txt”. The reads counts were normalized to TPM (Transcripts Per Million)^77^, which is a measurement of transcription level. DESeq2^78^ was used to calculate the differential expression genes, and the significance was determined by “fold change > 2 and FDR < 0.05”. Gene set enrichment analysis for gene ontology biological processes was used to analyze the genes of significant differential expression or genes fall into each TAD group or specific genomic regions using clusterProfiler R package^79^. All human genes were selected as background genes and only top significant terms were shown. The enrichments that passed the cutoff of FDR < 0.05 were considered significant.

### Assay for Transposase-Accessible Chromatin with high-throughout sequencing (ATAC-seq)

ATAC-seq was performed as described before with minor changes^80^. In brief, ∼10,000 cells were lysed in the lysis buffer (10 mM Tris-HCl (pH 7.4), 10 mM NaCl, 3 mM MgCl_2_ and NP-40) for 5 min on ice to extract the nuclei. After lysis, nuclei were spun at 500g for 5 min to remove the supernatant. Nuclei were then incubated with the Tn5 transposome and tagmentation buffer at 37 °C for 30 min (Vazyme, Cat. # TD502). After tagmentation, the stop buffer was added directly into the reaction to quench the tagmentation. DNA were cleaned by DNA Clean Beads, followed by a subsequent PCR step with 13 amplification cycles for adding additional index sequence to the fragments. The library were quantified by qPCR and the quality was assessed by Polyacrylamide Gel Electrophoresis (PAGE). Two biological replicates per condition were sequenced in a HiSeq X-Ten (PE150, Illumina) by Novogene (Beijing).

### ATAC-seq data analysis

To remove the adaptor of ATAC-seq paired data, the trim_galore software was performed with default parameters. Trimmed reads were mapped to human hg19 version reference genome using bowtie2 and converted to bam files using samtools^81^. ATAC peaks were identified by the bam files using MACS2^74^ with parameters -q 0.05 -- nomodel --shift -100 --extsize 200 for each sample. To annotate the peaks, the R package ChIPseeker was performed^82^. Then we used previously study^83^ to compare the accessibility between Pro-E and Ortho-E. Peaks were classified into different groups by the order of accessibility and the motifs of potential factor binding were enriched for each group. Motifs were identified using the “findMotifs” command of Homer^61^. The motifs with significant enrichment were selected to calculate the enrichment score, which was used to plot heatmap through ggplot2.

### Micrococcal nuclease (MNase)-seq

MNase-seq was performed as described before with minor changes^40^. Half a million FACS-purified primary erythroblasts were used for each reaction. Briefly, cells were lysed in MD buffer (50 mM Tris-HCl, 2 mM CaCl_2_, 0.2 % Triton X-100) supplied with 40U MNase for 5 min at 37°C to digest the chromatin, then 10µl STOP buffer (110 mM Tris-HCl, 55 mM EDTA) was added. After digestion with RNase A and Proteinase K, DNA fragments of 50bp-450bp was extracted for sequential library construction using the DNA library preparation kit (Vazyme, Cat. # ND607) following the manufacturer’s protocol. Two biological replicates per condition were sequenced in a HiSeq X-Ten (PE150, Illumina) by Novogene (Beijing).

### MNase-seq data analysis

MNase-seq reads were aligned to the human hg19 genome using bowtie2^72^ with default parameters after trimming the barcode. Only the reads with MAPQ > 30 and a unique match to the genome were kept for further analyses. The mitochondrial chromatin was also removed from analysis. In order to increase the resolution of nucleosome repeat length, we merged the replicates of each sample before progressing to further analysis. The nucleosome repeat length was calculated using Nuctools^84^. The CAM software^85^ was applied to determine the nucleosome position at regions around CTCF binging sites (2kb around weakened CTCF sites) derived from the CUT&RUN results.

### *In situ* Hi-C followed by chromatin immunoprecipitation (HiChIP) assay

HiChIP assay was carried out using the in-situ method as described before with minor modifications^33^. For GATA1 or KLF1 HiChIP, ten million Day-12 erythroblasts generated from ex vivo CD34+ erythroid differentiation culture were used; for H3K9me3 and H3K27ac HiChIP, one million sorted Pro-E or Ortho-E cells were used. In brief, after fixation with 1% formaldehyde, cells were digested for 2 hours at 37°C using 375U of MboI (NEB, Cat. #R0147). After biotin filling, proximity ligation was carried out with 4000U T4 DNA Ligase (Vazyme, Cat. # N103-01) for 4 hours at 25°C. Sonication of re-suspended nuclei was performed by Covaris M220 sonicator. Following an overnight incubation with specific antibodies, 50µl Protein A/G Magnetic Beads were added at 4°C and incubated for 2 hours. Bound DNA–protein complexes were eluted and reverse-crosslinked after a series of washes.

After reverse-crosslinking, ligation fragments containing biotin were immobilized on MyOne Streptavidin C1 beads (ThermoFisher Cat.#: 65602), and the library was prepared using the Tn5 transposome DNA library preparation kit (Vazyme, Cat.# TD502) as per the manufacturer’s protocol. DNA was then subjected to double-size selection using DNA Clean Beads (Vazyme, Cat. # N411) in order to isolate fragments between 300 and 600bp. Two biological replicates per condition were sequenced in a HiSeq X-Ten (PE150, Illumina) or NovaSeq 6000 (S4, PE150, Illumina) by Novogene (Beijing).

### HiChIP data analysis

HiChIP sequence reads were mapped to the human reference genome hg19 using HiC-Pro (version 2.10.0)^59^. HiC-Pro output results were used as input of hichipper to call high-confidence chromatin loops through “EACH, ALL” parameter settings^86^. To visualize loop interactions identified by hichipper, the interaction files compatible with the WashU browser were generated through adding “-mw” parameter. HiChIP loops of GATA1 and KLF1 were filtered to meet the criteria of min/max (5kb/2Mb) distance as well as False Discovery Rate (FDR) < 0.05. H3K9me3 HiChIP loop length was more than 5kb with FDR < 0.05. Then we quantified the loop length percentage using the data after filtering. To illustrate the increase of B-B interactions and reductions of A-A interactions in Ortho-E, H3K9me3 associated interactions were used to calculate intra-compartment and inter-compartment interaction frequency.

### Immunoblotting

Whole-cell lysates preparation and Western blot analyses were carried out as described previously^28^. In brief, cells were harvested, washed in ice-cold PBS, and the protein concentration of the extracts was determined using bicinchoninic acid reagent (Thermo Fisher Scientific). Equal amount of protein (10 µg per lane) was loaded, separated using 15% SDS-PAGE gel and transferred onto PVDF membranes. The membranes were blocked with 5% non-fat milk, and then incubated with the primary antibodies at 4 °C overnight. Subsequent to incubation with the respective secondary antibody, immune complexes were detected using ECL plus Western blotting reagents (Thermo Fisher Scientific). The expression levels of β-actin or total histone H3 were monitored as an internal control. Total protein level of each lane was normalized by VersaBlot™ Total Protein Normalization Kits (Biotium Cat. # 33025).

### Immunofluorescence staining and imaging analyses

FACS-sorted cells were rinsed three times with PBS, fixed with 4% paraformaldehyde, and then permeabilized with 0.1% Triton X-100. To eliminate the interference from membrane-bound fluorescence-labeled FACS antibodies, M.O.M Kit (Vector) was used to block the FACS antibodies (Mouse IgG). Cells were incubated with the primary antibody at 4℃ overnight, washed and then incubated for 2 hours with the secondary antibody at room temperature. The samples were stained with DAPI and then mounted with ProLong Glass reagent (Thermo Fisher Scientific).

Nikon A1RS+ confocal microscope was used to acquire images of stained cells (objective lens used: CFI Apochromat TIRF 100XC Oil (NA 1.49)). The images were analyzed using NIS Elements software, and the z stack of optical sections was used to generate 3D reconstruction.

## Data availability

Data in this manuscript have been deposited in the Genome Sequence Archive (GSA) in the national genomics data center. The assigned accession of the submission is HRA001026. The code is available upon request. The supplement materials can be found in Supplementary Tables 1-7.

## Acknowledgments

We thank the Flow Cytometry Core and the Imaging Core of the National Center for Protein Sciences at Peking University (PKU). We are grateful to Dr. Hongxia Lyu, Fei Wang, Yinghua Guo, Dr. Chunyan Shan for the technical support. We thank Dr. Zheng Wang at CAMS&PUMC for the help with immunofluorescence assay and proofreading of the manuscript. We thank Dr. Xiong Ji, Dr. Yongpeng Jiang and Rui Wang at PKU for helping with the Hi-C method. We thank Dr. Cheng Li and Dr. Chao Zhang for the advice on Hi-C data analysis.

This project was supported by National Natural Science Foundation of China (81970110), by Peking-Tsinghua Center for Life Sciences and College of life science, Peking University to H.Y.L.. This work was supported by the National Key Research and Development Program (2017YFA0104500), the Foundation for Innovative Research Groups of the National Natural Science Foundation of China (81621001) to X.J.H..

## Author contributions

D.L., F.W., X.J.H. and H.Y.L. conceptualized this project and designed the experiments. D.L. and S.Z. performed the experiments. F.W. and D.L. conducted bioinformatic analyses. D.L., F.W., X.J.H. and H.Y.L. wrote the manuscript. All authors discussed the results and commented on the manuscript.

---

**Extended Data Fig. 1 (Fig 1):**
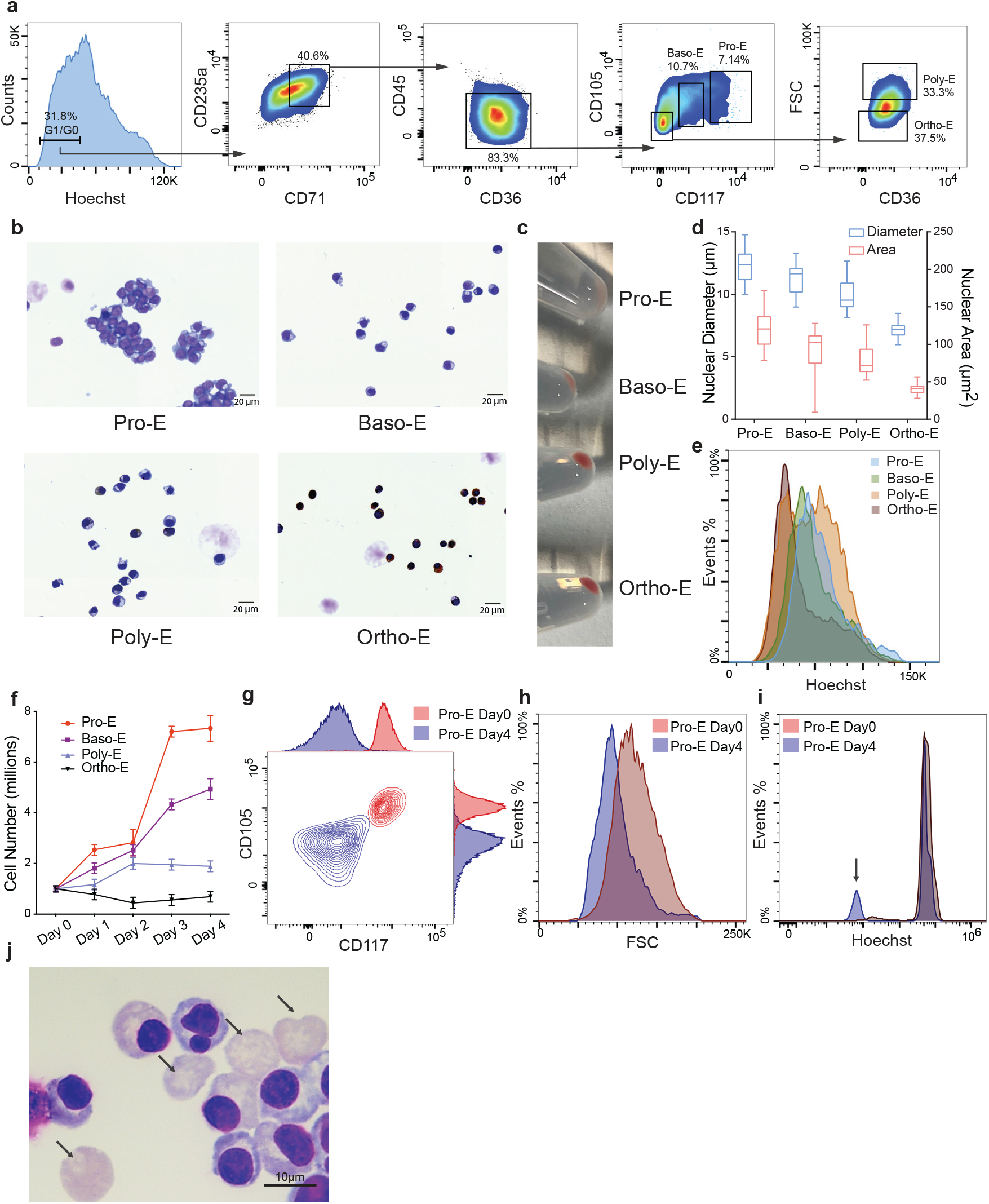
**a**, Fluorescence-activated cell sorting (FACS) scheme for isolating primary human terminal erythroblasts of distinct stages from the *ex vivo* CD34^+^ erythroid differentiation system. To avoid the effects of cell cycle, G0/G1 cells are gated based on the signal of Hoechst33342. CD235a^+^ CD71^+^ CD45^low^ CD36^+^ CD117^high^ CD105^high^ cells are pro-erythroblast cells (Pro-E), CD235a^+^ CD71^+^ CD45^low^ CD36^+^ CD117^dim^ CD105^dim^ cells are basophilic erythroblasts (Baso-E), CD235a^+^ CD71^+^ CD45^low^ CD36^+^ CD117^-^ CD105^-^FSC^high^ cells are polychromatic erythroblasts (Poly-E), and CD235a^+^ CD71^+^ CD45^low^ CD36^+^ CD117^-^ CD105^-^ FSC^low^ cells are orthochromatic erythroblasts (Ortho-E). **b**, Representative images of purified primary Pro-E, Baso-E, Poly-E and Ortho-E cells by benzidine-Giemsa staining. **c**, Pellets of 1 million cells at the indicated stages demonstrating accumulation of hemoglobin along with erythroid differentiation. **d**, Nuclear diameter and area of sorted Pro-E, Baso-E, Poly-E and Ortho-E cells. (N = 200) **e**, Cell cycle profiles of Pro-E, Baso-E, Poly-E and Ortho-E cells without gating for specific cell cycle stages. **f**, Proliferation assay of FACS-purified Pro-E, Baso-E, Poly-E and Ortho-E cells. (Starting from 1 million cells/sample, N = 4) **g**, Expression of CD105 and CD117(c-Kit) on cell surface of uncultured (Day 0) and four-day cultured (Day 4) Pro-E cells. **h**, Cell size (indicated by forward scatter, FSC) of uncultured (Day 0) and four-day cultured (Day 4) Pro-E cells. **i**, Hoechst33342 staining of uncultured (Day 0) and four-day cultured (Day 4) Pro-E cells. The arrow indicates enucleated reticulocytes. **j**, Representative image of four-day cultured (Day 4) Pro-E cells by Giemsa staining. (Arrows indicate enucleated reticulocytes.)

**Extended Data Fig. 2 (Fig 1):**
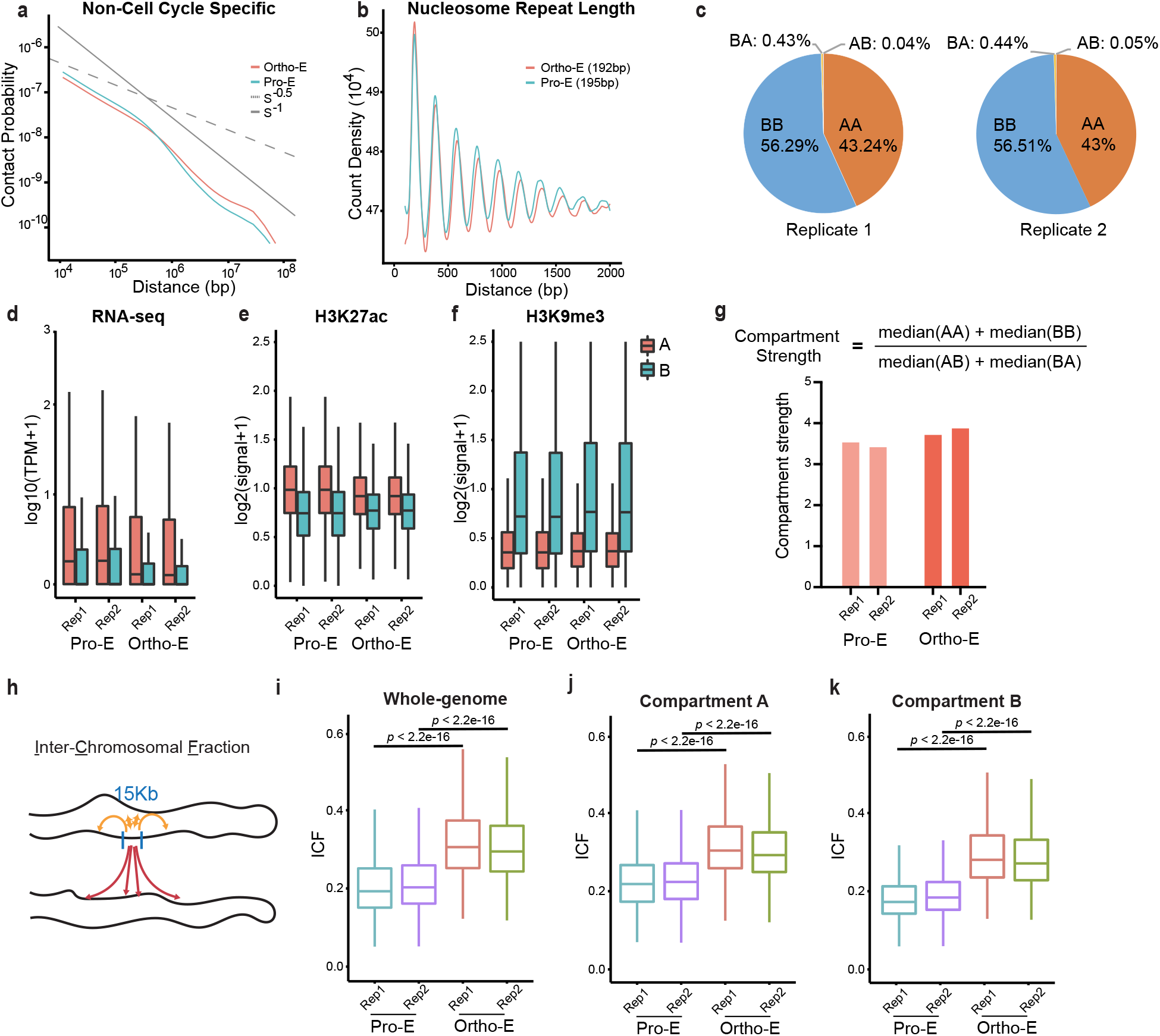
**a**, Hi-C contact probability as a function of distance of non-cell cycle specific Pro-E and Ortho-E. P(s)∼S^-0.5^ indicates the mitotic state, ∼S^-1^ indicates the fractal globule state. **b**, Estimation of NRL (nucleosomal-repeat length) based on MNase-seq in Pro-E and Ortho-E cells. **c**, Compartment analysis in Pro-E and Ortho-E based on PC1 value of Hi-C assay. (AA: compartment A in Pro-E, which remains as compartment A in Ortho-E; BB: compartment B in Pro-E, which remains as compartment B in Ortho-E; AB: compartment A in Pro-E, which switches into compartment B in Ortho-E; BA: compartment B in Pro-E, which switches into compartment A in Ortho-E) **d**, Normalized signal of RNA-seq in compartment A or B of Pro-E and Ortho-E. TPM, Transcripts Per Kilobase Million. **e**, Normalized H3K27ac signal by CUT&RUN assay in compartment A or B of Pro-E and Ortho-E. **f**, Normalized H3K9me3 signal by CUT&RUN assay in compartment A or B of Pro-E and Ortho-E. **g**, (Top) Formula for quantification of compartment strength; (bottom) compartment strength in Pro-E and Ortho-E. **h**, Schematic description of ICF (Inter-Chromosomal Fraction). **i**, ICF at whole-genome level of Pro-E and Ortho-E. **j**, ICF in compartment A of Pro-E and Ortho-E. **k**, ICF in compartment B of Pro-E and Ortho-E.

**Extended Data Fig. 3 (Fig 2):**
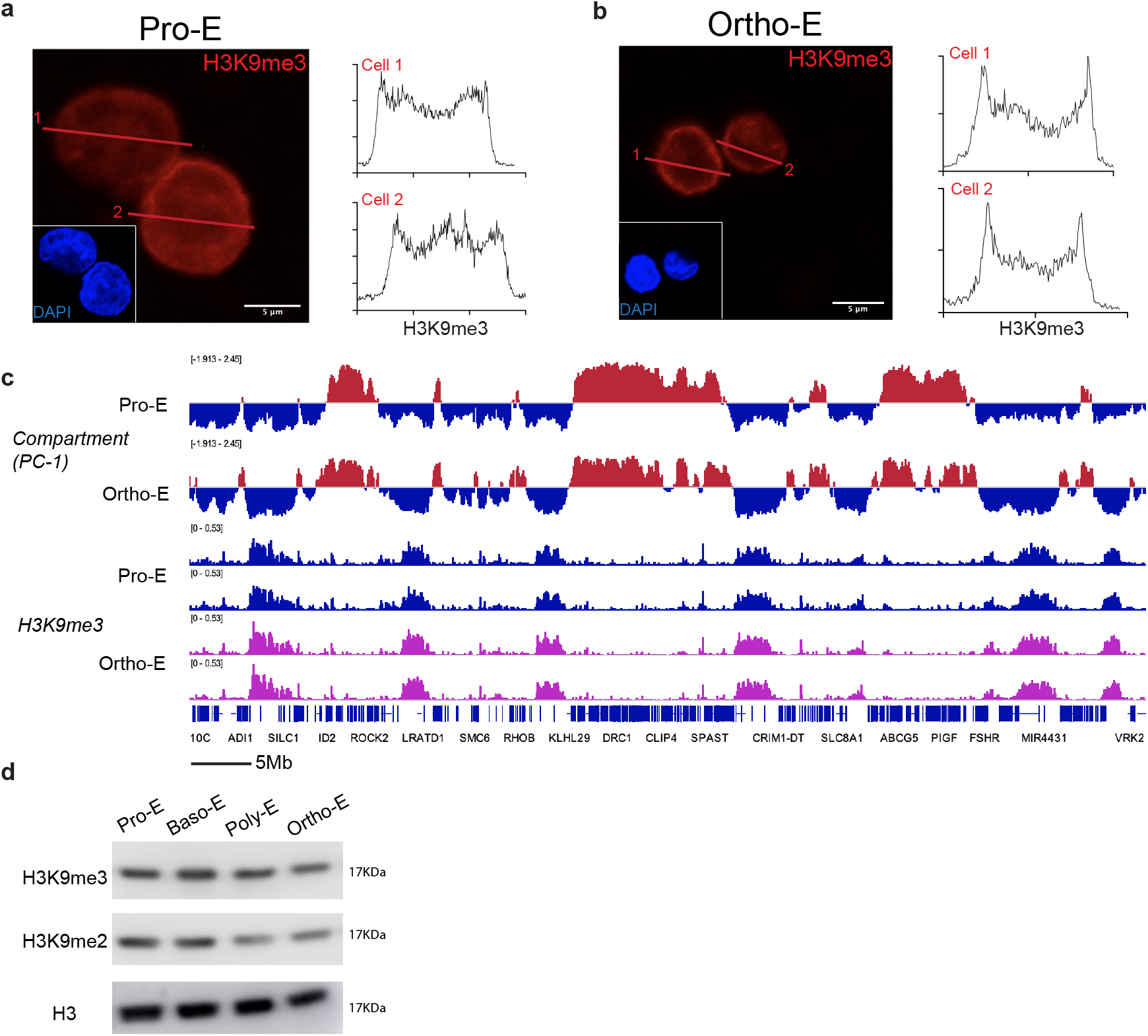
**a**, (Left) Immunofluorescence images of heterochromatin marker H3K9me3 (red) in Pro-E cells; DNA was counterstained by DAPI (blue). (Right) Signal distribution of H3K9me3 across the corresponding line of cells indicated. **b**, (Left) Immunofluorescence images of heterochromatin marker H3K9me3 (red) in Ortho-E cells; DNA was counterstained by DAPI (blue). (Right) Signal distribution of H3K9me3 across the corresponding line of cells indicated. **c**, Compartment (PC1 value of Hi-C) and H3K9me3 (CUT&RUN) distribution of Pro-E and Ortho-E at Chr2: 0-60Mb. Red: compartment A; blue: compartment B. **d**, Immunoblotting of H3K9me3 and H3K9me2 in Pro-E, Baso-E, Poly-E and Ortho-E cells.

**Extended Data Fig. 4 (Fig 3):**
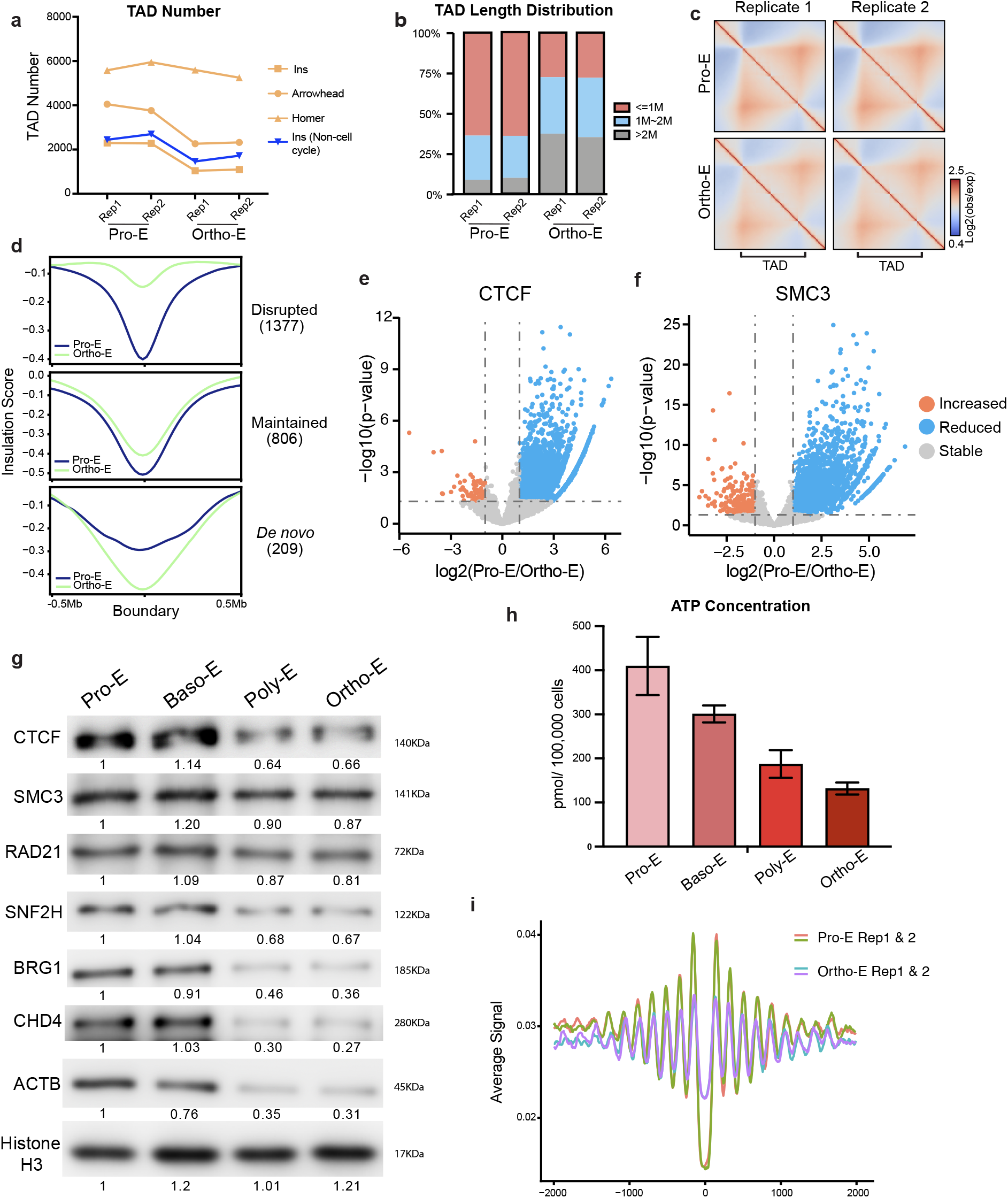
**a**, TAD numbers called by different algorithm in Pro-E and Ortho-E cells. G0/G1 samples are used for analysis unless noted. **b**, TAD length distribution in Pro-E and Ortho-E. **c**, Heatmaps showing the normalized average interaction frequencies for all TADs (defined in Pro-E) as well as their nearby regions (±0.5 TAD length) in Pro-E and Ortho-E. **d**, Average insulation score of the Disrupted, Maintained and De novo formed TAD boundaries in Pro-E and Ortho-E. **e**, Volcano plot of differential and constant binding of CTCF between Pro-E and Ortho-E. **f**, Volcano plot of differential and constant binding of SMC3 between Pro-E and Ortho-E. **g**, Immunoblotting of CTCF, SMC3, RAD21, SNF2H, BRG1, CHD4, ACTB and Histone H3 in Pro-E, Baso-E, Poly-E and Ortho-E stages. (Quantification of protein level was based on total protein staining for each sample using the entire lane.) **h**, Quantification of ATP level in Pro-E, Baso-E, Poly-E and Ortho-E cells. **i**, Nucleosomal signal (based on MNase-seq) at the sites with CTCF loss from Pro-E to Ortho-E.

**Extended Data Fig. 5 (Fig 4):**
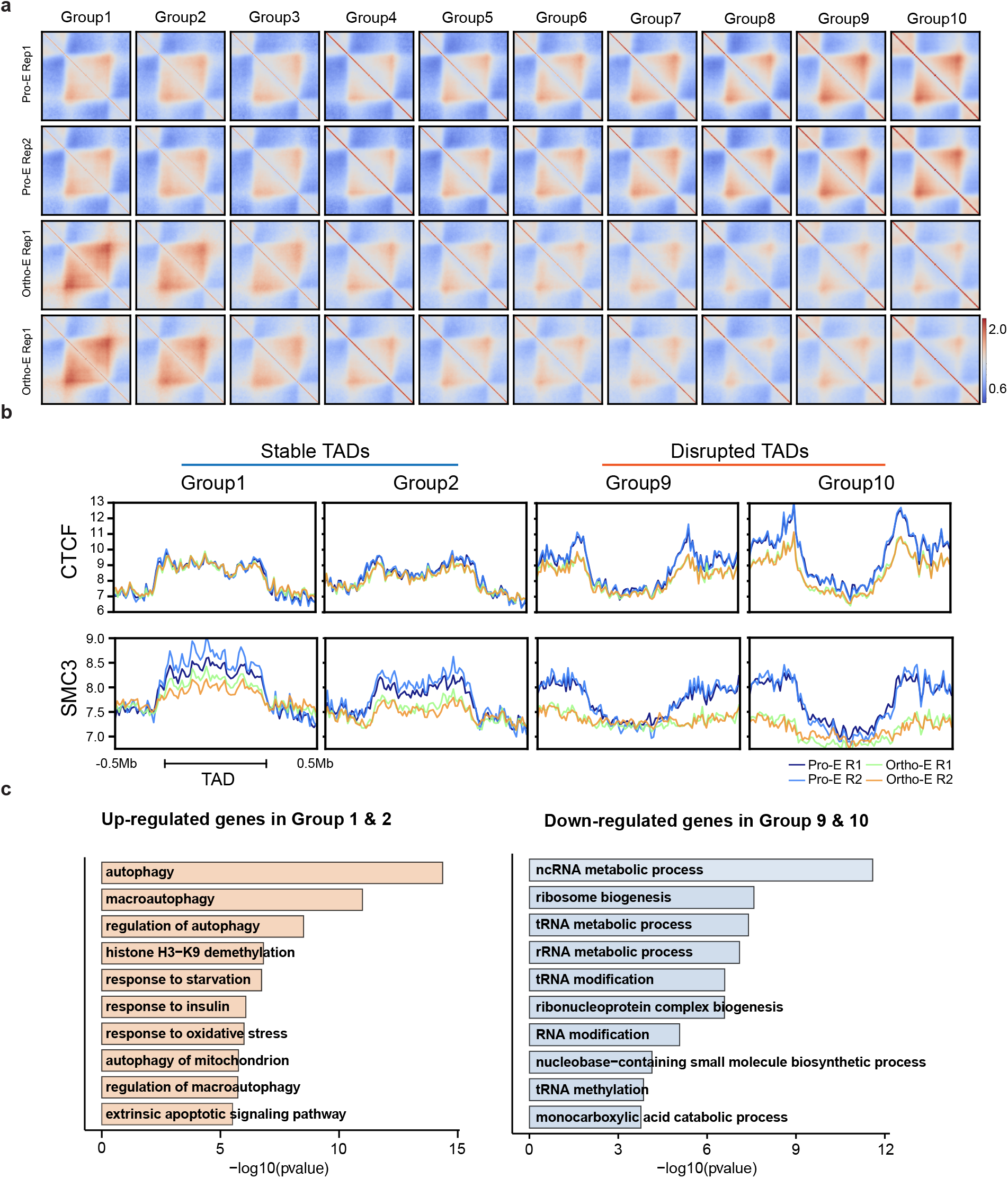
**a**, Heatmaps showing the normalized average interaction frequencies for TADs and the neighboring ±0.5TAD regions. The TADs are grouped by the TAD score ratio (TAD score of Ortho-E to that of Pro-E). Group1 are the most “Stable” TADs; the stability of TADs goes down gradually from group 1 through group 10. **b**, Distribution of CTCF and SMC3 signal by CUT&RUN assay in TADs and the neighboring ±0.5 Mb regions. The TADs are grouped by their TAD score ratio. Group1&2 represent the most “Stable” TADs; group 9&10 represent the most “Disrupted” TADs. **c**, (Left) Top ten GO terms of up-regulated genes in TAD regions of Group 1&2; (right) top ten GO terms of down-regulated genes in TAD regions of Group 9&10.

**Extended Data Fig. 6 (Fig 5):**
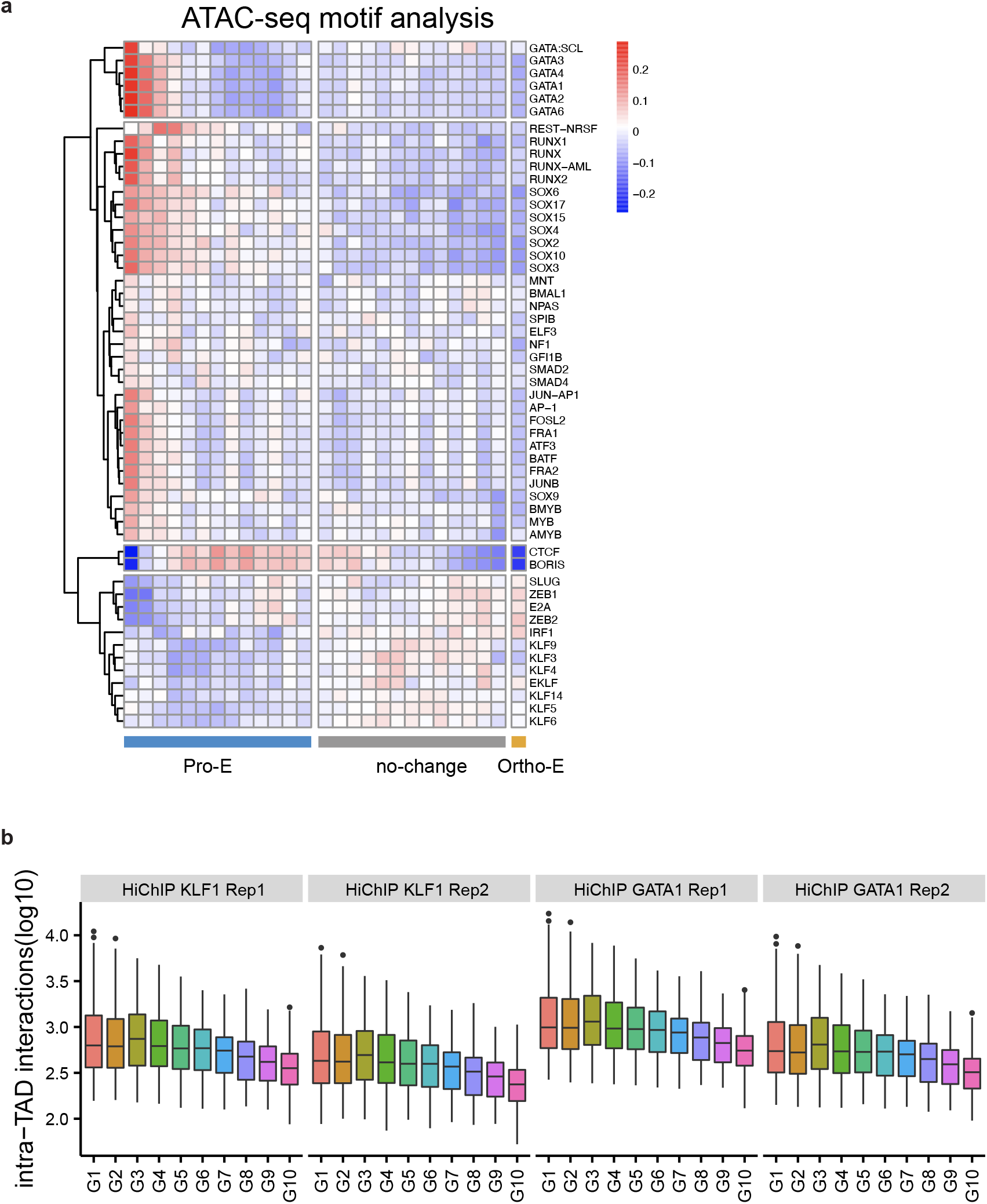
**a**, Heatmap showing the motif enrichment score based on ATAC-seq assay. Peaks of differential chromatin accessibility in ProE vs. Ortho-E were ranked by the fold change, and then grouped into ∼1500 peaks/bin. (Each bin is shown as a square in the plot.) The bins from left to right were arranged from the chromatin accessible sites that are most enriched in Pro-E to the ones that are most enriched in Ortho-E. *De novo* motif search was conducted in the peaks of the same bin. The factors of enriched motifs are listed on the right of the plot. **b**, GATA1 or KLF1 mediated intra-TAD interaction frequency within TAD groups 1∼10.

**Extended Data Fig. 7 (Fig 6):**
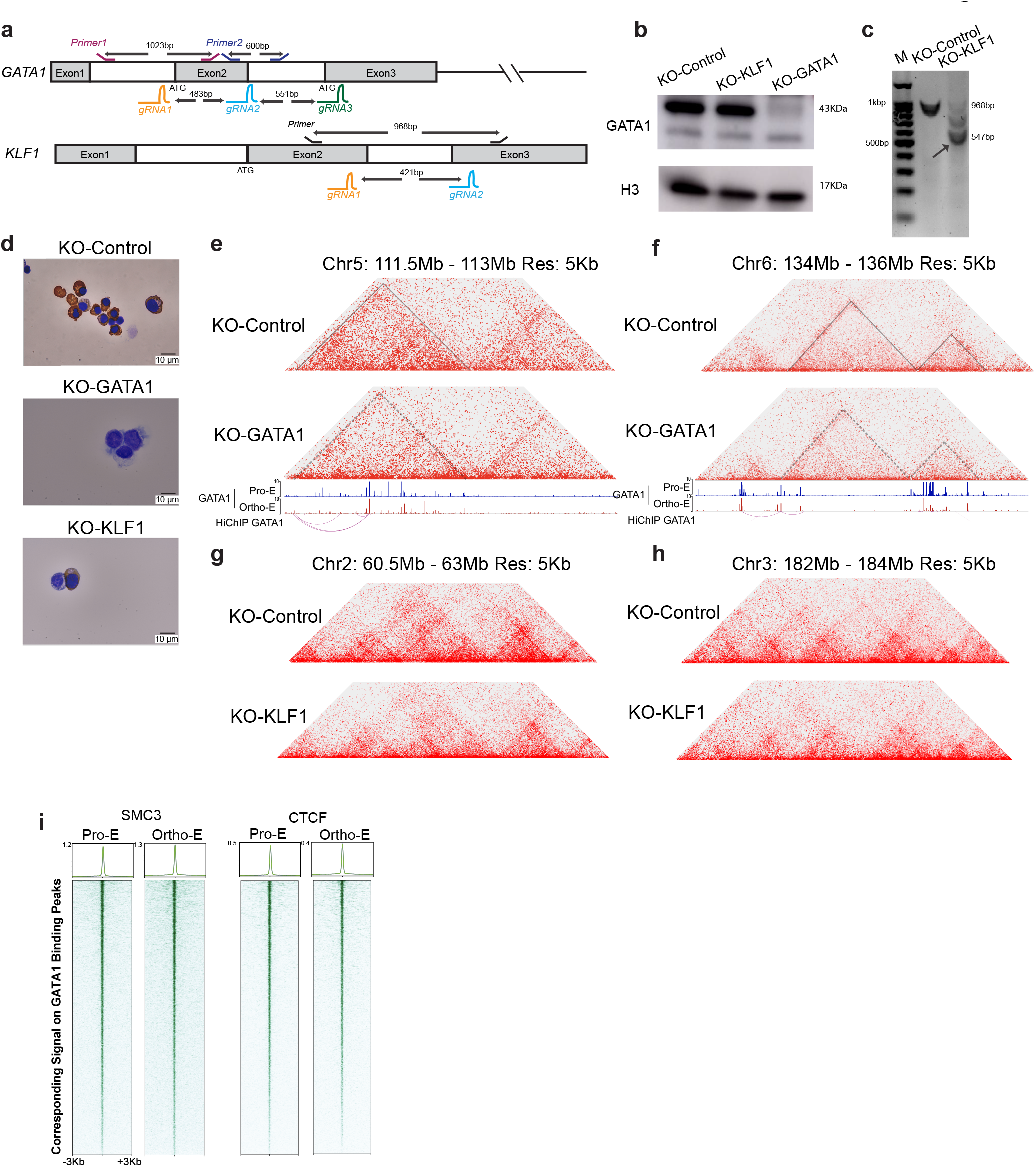
**a**, Schematic description of GATA1 and KLF1 knockout strategies using CRISPR/Cas9 in purified primary human Pro-E cells. **b**, Immunoblotting of GATA1 after knockout by control, KLF1 or GATA1 sgRNA for 48 h in Pro-E. Histone H3 was shown as the loading control. **c**, Genotyping of KLF1 loci after knockout with control or KLF1 sgRNA for 48 h. (The arrow indicates the re-joined DNA after complete fragment removement by paired gRNAs.) **d**, Images of Pro-Es after knockout by control, GATA1 or KLF1 sgRNA for 72 h by benzidine-Giemsa staining. **e&f**, (Top) Hi-C contact matrices of Pro-Es after knockout by control or GATA1 sgRNA for 72 h at chromosome regions indicated; (middle & bottom) corresponding CUT&RUN and HiChIP signal of GATA1. Juicebox parameters were balanced normalized at 5 kb resolution. Dashed triangle shows the disrupted TAD resulting from knocking-out of GATA1. **g&h**, Hi-C contact matrices of Pro-Es after knockout by control or KLF1 sgRNA for 72 h at chromosome regions indicated. Juicebox parameters were balanced normalized at 5 kb resolution. **i**, Density maps showing corresponding SMC3 and CTCF CUT&RUN signals at the GATA1 peaks in Pro-E and Ortho-E cells.

**Extended Data Fig. 8:**
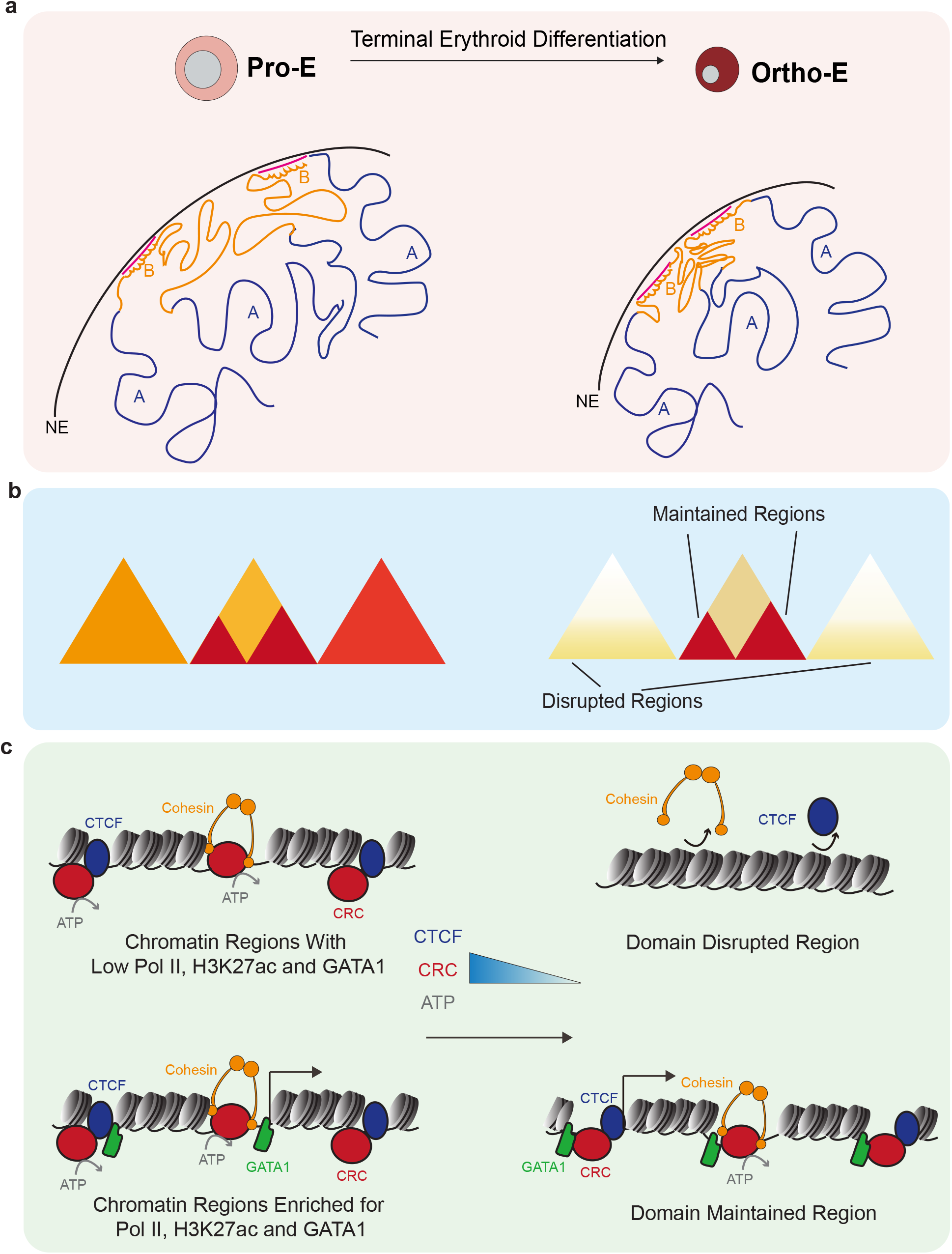
Model of chromatin compaction and domain disruption in terminal erythropoiesis. During terminal erythroid differentiation, the chromatin is rearranged at multi-dimensional level to achieve nuclear compaction to 20-30% of the original volume. **a**, heterochromatin reorganizes via establishing long-range interactions and aggregating at the sub-nuclear membrane region. (NE: Nuclear envelope; A: Compartment A; B: Compartment B). **b**, a large number of TADs undergo disruption, while select TADs enriched for transcription competence marks are maintained until the end of erythroid differentiation. **c**, the early erythroblasts (Pro-E) contain sufficient energy to sustain the nuclear structure relapse between normal cell cycles; the level of ATP, CTCF and chromatin remodeling complex (CRC) is reduced as terminal differentiation progresses, and therefore, the late erythroblasts (Ortho-E) can only maintain the chromatin architecture at select essential regions, which are enriched for transcription activity and/ or critical transcription factor binding such as GATA1.

